# Marsupial immune protection is shaped by enhancer sharing and gene cluster duplication of cathelicidin antimicrobial peptides

**DOI:** 10.1101/2024.07.29.605640

**Authors:** Jongbeom Park, Wenfan Ke, Aellah Kaage, Charles Y. Feigin, Yuri Pritykin, Mohamed S. Donia, Ricardo Mallarino

## Abstract

Marsupial neonates are born with immature immune systems, making them vulnerable to pathogens. While neonates receive maternal protection, they can also independently combat pathogens, though the mechanisms remain unknown. Using the sugar glider (*Petaurus breviceps*) as a model, we investigated immunological defense strategies of marsupial neonates. Cathelicidins – a family of antimicrobial peptides expanded in the genomes of marsupials – are highly expressed in developing neutrophils. Sugar glider cathelicidins reside in two genomic clusters and their coordinated expression is achieved by enhancer sharing within clusters and long-range physical interactions between clusters. These cathelicidins modulate immune responses and have potent antimicrobial effects, sufficient to provide protection in a mouse model of sepsis. Lastly, cathelicidins have a complex evolutionary history, where marsupials and monotremes are the only tetrapods that retained two cathelicidin clusters. Thus, cathelicidins are critical mediators of marsupial immunity, and their evolution reflects the life history-specific immunological needs of these animals.

## Introduction

Prenatally protected in the sterile maternal womb, mammalian offspring are suddenly exposed to a plethora of microbes at birth (*1–3*). While such exposure allows colonization of commensal microbes, it also poses a significant threat, as sepsis caused by infections can lead to rapid neonatal mortality (*4–6*). Notably, susceptibility to such infections may vary among different mammalian lineages as a function of their life history. For instance, precocial mammals, such as cattle and guinea pigs, are often equipped with most of the essential immune components at birth (*7*, *8*). This relative immunological maturity at birth enables precocial mammals to swiftly respond to pathogens. In contrast, altricial mammals, such as mice and rats, are born with a less developed immune system due to their short gestation period (*9*, *10*), requiring more extensive maternal care compared to their precocial relatives.

Among mammals, marsupials are an extreme case of altricial birth. Marsupials diverged from eutherians around 160 million years ago and constitute a unique lineage with characteristic reproductive and morphological traits (*11*, *12*). Females have short pregnancies and give birth to highly immature young that reside inside a pouch, where they complete their physical development. A consequence of this short gestation period is that several key immune components are absent in marsupial neonates. Namely, while the hematopoietic niche has already migrated from the liver to the bone marrow during eutherian fetal development, the marsupial neonate liver is still an active site of hematopoiesis (*13*, *14*). Moreover, unlike eutherian neonates, marsupial newborns lack lymphoid organs such as thymus and lymph nodes, making them incapable of mounting an adaptive immune response. This presents a significant challenge because the maternal pouch is in contact with the environment and is known to harbor a wide range of bacterial species, including many pathogenic ones (*15*, *16*).

Despite their immunological immaturity, marsupial neonates can survive the non-sterile environment of the pouch in part through maternal protection. This includes the transfer of immunoglobulins through milk as well as the secretion of antimicrobial compounds from specialized pouch glands (*17–21*). Additionally, various immune-related tissues in developing marsupials, including bone marrow and spleen, contain granulocytes and lymphocytes, indicating that neonates themselves can actively fend off pathogen attacks (*13*). Our current understanding of marsupial neonatal immune protection, particularly neonate-mediated immunity, remains incomplete for several reasons. First, previous research on marsupial immunity has largely relied on work conducted in wild animals (*18*, *22–24*), making it challenging to obtain accurately timed samples or perform controlled experiments. Additionally, the scarcity of suitable antibodies has hindered efforts to identify cell types and the cellular composition of marsupial neonatal hematopoietic tissue using histology and microscopy (*13*, *25–30*). Finally, to our knowledge, there has been no unbiased experiment conducted to identify key protective genes expressed by neonatal immune cells. Here, using our captive colony of marsupial sugar gliders (*Petaurus breviceps*) (Fig. 1A) and combining transcriptomics, epigenomics, functional assays, and comparative genomics, we set out to dissect the mechanisms underlying neonate immune protection in marsupials.

**Figure 1.**
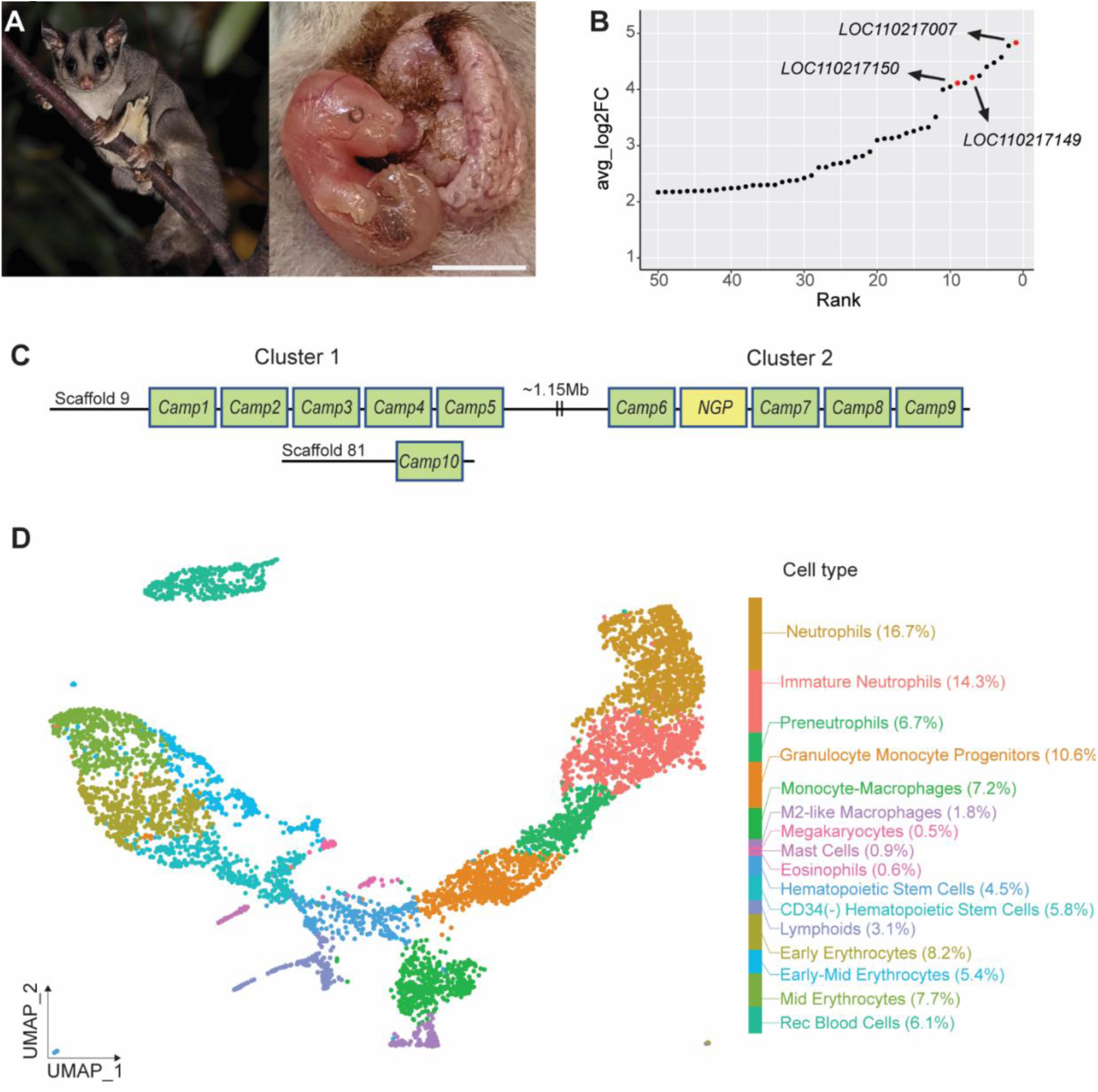
Cathelicidins are highly and specifically expressed in sugar glider neutrophils. (**A**) An adult sugar glider (left panel; photo by Patrick Kavanagh) and a postnatal day 4 joey laying atop the everted maternal pouch (right panel). Scale bar: 0.5cm. (**B**) Graphical representation of the top differentially expressed genes in neutrophil lineage cells compared to the rest of the liver cells. Genes were ranked by log2FC values. Cathelicidin genes are denoted in red. (**C**) A schematic of the genome structure of sugar glider cathelicidins. Reciprocal blast with leadbeater’s possum suggests that *Camp10* belongs to Cluster 1. (D) UMAP plot showing cell type clustering and composition of the sugar glider neonatal (P0) liver.

## Results

### Neutrophils are the most abundant immune cells in the sugar glider neonate liver

While histological studies have identified few immune cells in developing marsupial neonates, the exact composition and identity of these immune cells during hematopoiesis, as well as the repertoire of key protective genes expressed by each cell type, are unknown. To fill this gap, we conducted single-cell RNA-seq (scRNA-seq) of the day of birth (postnatal day 0, P0) sugar glider liver, which is the main site of neonatal hematopoiesis at this stage (*13*). We detected 6028 cells grouped into 16 clusters, which we then annotated using established gene expression markers (fig. S1). Our analysis revealed that neutrophil lineage cells (i.e., neutrophils, immature neutrophils, preneutrophils, and granulocyte monocyte progenitors) were the most abundant cell type, comprising 48.4% of the entire cell population. Neutrophils are a type of white blood cell and are an essential part of the innate immune system, the body’s first line of defense against infections (*31*). Red blood lineage cells were the second most abundant cell type, comprising 27.4% (fig. S1). Interestingly, we identified a small lymphoid cell population (3.2% of cells), a finding that contrasts with previous reports suggesting that T and B cells were not present in the liver of marsupial neonates (fig. S1) (*13*, *14*, *32*, *33*). Thus, by characterizing the cell type composition of the marsupial liver at single-cell resolution, our results suggest that marsupials rely heavily on neutrophils for neonatal protection.

### Cathelicidins are highly expressed in marsupial neutrophils

The finding that developing neutrophils are the most abundant cells in marsupial neonatal hematopoietic tissue prompted us to focus on this lineage and investigate its function. We therefore conducted differential expression analyses comparing all neutrophil lineage cells to other cell types in the liver. Among the 596 genes that were differentially expressed (P-value < 0.0001, Log2Fold > 1), three of the top ten most highly upregulated genes in neutrophil lineage cells (*LOC11021700, LOC110217149*, and *LOC110217150*), including the most highly upregulated one (*LOC11021700*), were identified as cathelicidin antimicrobial peptides (AMPs) (Fig. 1B and data table S1). Cathelicidins constitute a key component of the innate immune system due to their ability to disrupt microbial membranes and modulate immune responses (*34–36*).

Notably, compared to humans and mice, marsupial genomes have a much larger repertoire of cathelicidin genes. The peptides translated from these genes are known to play key roles in regulating immune responses of different marsupial species (*22*, *23*, *37–43*). Thus, to gain more insights into the evolution and function of marsupial cathelicidins, we next characterized the expression of these genes in sugar gliders. To do this, we first systemically annotated all members of this gene family in the sugar glider genome, then performed expression-based validation (see materials and methods) (table S1 and data table S2) (*44*). In total, we identified ten cathelicidin antimicrobial peptide genes, labeled *Camp1-10*, as well as the sugar glider ortholog of *Ngp* (neutrophilic granule protein), a cathelicidin-related gene. These genes resided in two clusters located 1.15Mb apart and distinguished by the phylogenetic affinity of their constituent genes (Fig. 1C, figs. S2 and S3, and table S2). Cluster 1 contains *Camp1-Camp5* and *Camp10,* whereas Cluster 2 contains *Camp6-9* and *Ngp*. While *Camp2* and *Camp5* have stop codons in exon 2, indicating that they are likely pseudogenes, the remaining genes encode proteins that range from 100-266 amino acids in length and 11.1-70.9% in identity (table S3).

Next, we reanalyzed our scRNA-seq data from neonatal liver to establish the specific cell types in which the different cathelicidins and *Ngp* are expressed (Fig. 1D). Most of these genes (i.e., *Camp1, Camp3*, *Camp4*, *Camp6, Camp8*, *Camp9*, *Camp10*, and *Ngp*) are expressed exclusively in neutrophil lineage cells (Fig. 2A and fig. S4). Among neutrophil lineage cells, we observed differences in expression levels, with *Camp3*, *Camp4*, *Camp10*, *Camp6*, and *Ngp* showing high, overlapping expression levels in a large proportion of cells, whereas *Camp1*, *Camp8*, and *Camp9* are expressed at relatively lower levels and in a reduced number of cells (Fig. 2A and figs. S4 and S5). Interestingly, *Camp7* is exclusively expressed by M2-like macrophage cells, suggesting that its transcriptomic regulation, and likely its function, are distinct from the other cathelicidin gene family members (Fig. 2A and fig. S5).

**Figure 2.**
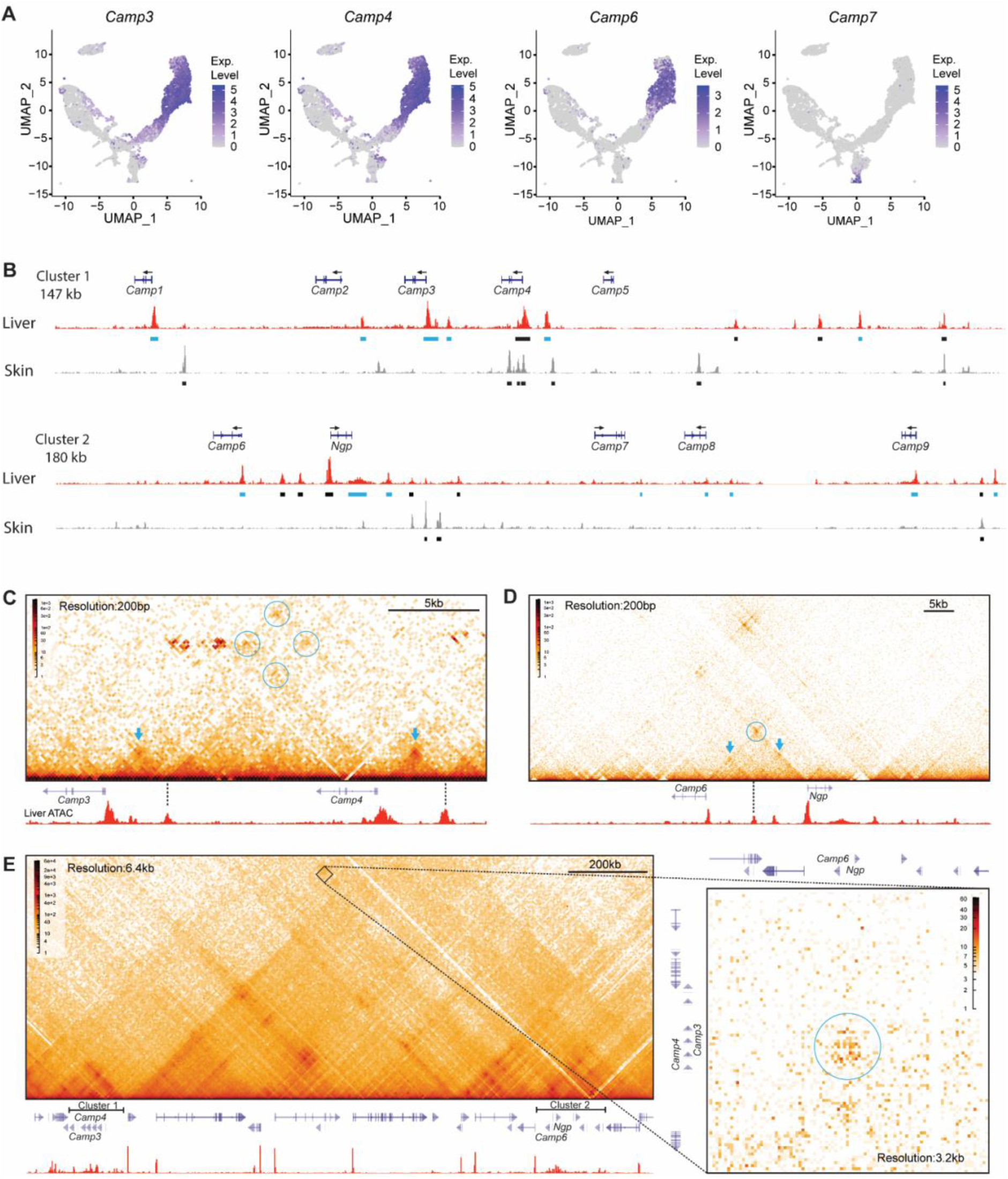
Expression and regulation of sugar glider cathelicidins. (**A**) UMAP plots displaying the expression of selected cathelicidin genes. (**B**) ATAC-seq traces of sugar glider liver and skin tissue. Shown are genomic regions corresponding to clusters 1 and 2. Black boxes denote putative cis-regulatory elements (CREs) and blue boxes denote putative CREs containing *Cebp* family binding sites. Arrows indicate the direction of transcription. (**C-E**) Contact maps of sugar glider adult bone marrow. Displayed are genomic regions containing *Camp3* and *Camp4* (**C**), *Camp6* and *Ngp* (**D**), and both cathelicidin clusters (**E**). Heatmaps visually represent the frequency of physical interactions between different regions of the genome, with darker colors representing higher frequencies than lighter colors. Blue arrows denote an interaction between enhancers and their proximal promoters. Circles indicate interactions among multiple enhancers and promoters.

Finally, we characterized the dynamics of cathelicidin expression in hematopoietic tissues throughout the lifetime of sugar gliders by performing longitudinal bulk RNA-sequencing on liver tissue from P0, P10 and adult (> 2 years) sugar gliders. In addition, we sampled bone marrow from adult sugar gliders, as this tissue is the main site of hematopoiesis in adult marsupials (*13*). We found that five of the cathelicidins (*Camp3*, *Camp4*, *Camp10*, *Camp6*, and *Camp7*) as well as *Ngp* are expressed in moderate to high levels in the liver of P0 and P10 joeys, while their expression in this tissue is reduced in adults (Read per kilobase per million reads (RPKM)>50) (fig. S6). Notably, these same genes, as well as *Camp9*, are highly expressed in adult bone marrow (fig. S6).

Taken together, our transcriptomic analyses indicate that cathelicidins have divergent expression patterns, with a subset of them - *Camp3*, *Camp4*, *Camp10*, *Camp6*, and *Ngp* – showing high expression levels in overlapping neutrophil lineage cells, while the expression of another one – *Camp7* – is restricted to macrophage cells. Moreover, our longitudinal analysis indicates that these same genes continue to be expressed at high levels throughout the lifetime of sugar gliders, first in the liver and subsequently in the bone marrow.

### Co-expression of cathelicidins is driven by enhancer sharing

Considering that a subset of cathelicidin genes (i.e., *Camp3*, *Camp4*, *Camp6*, *Camp10*, and *Ngp*) were highly co-expressed in developing neutrophils, we next sought to determine whether these genes share a common regulatory apparatus. To this end, we conducted an assay for transposase-accessible chromatin (ATAC-seq) in P10 sugar glider liver, an approach that allows for the identification of open chromatin regions and constitutes a useful strategy for identifying putative cis-regulatory elements (CREs) (*45*, *46*). We identified a total of 10 and 14 peaks located in Clusters 1 and 2, respectively (Fig. 2B). To further filter our data, we reanalyzed ATAC-seq data from sugar glider skin tissue (*47*), which, with the exception of *Camp7*, does not express cathelicidin genes (*44*). Thus, we could exclude shared open chromatin regions between these tissues as plausible regulators of cathelicidin gene expression. Examination of candidate CREs showed that *Camp3*, *Camp4*, *Camp6*, and *Ngp*, all of which were highly expressed in liver neutrophils, had a total of seven liver-specific open peaks in nearby intergenic regions (Fig. 2B).

We next examined whether any of these putative CREs physically interact with the promoters of *Camp3*, *Camp4*, *Camp6*, and *Ngp* to regulate the expression of these genes. To achieve this, we sampled bone marrow tissue from adult sugar gliders and performed Region Capture Micro-C (RCMC), a chromosome conformation capture technique that enables the detection of interactions between genomic loci at high resolution (*48*). Among the putative CREs found in Cluster 1, two of them displayed strong contact interactions with the promoters of *Camp3* and *Camp4* (Fig 2C). Similarly, we identified one putative CRE in Cluster 2 showing a strong contact interaction with the promoters of *Camp6* and *Ngp* (Fig 2D). Remarkably, in addition to within cluster interactions, our RCMC data revealed a long-range (>1 Mb) contact interaction between clusters. Specifically, we found evidence of contacts between genes found in Cluster 1 (i.e., *Camp3* and *Camp4*) and genes found in Cluster 2 (i.e., *Camp6* and *Ngp*) (Fig 2E). This result suggests that genes across both clusters can be co-regulated.

Motivated by the finding that enhancer sharing can account for the coordinated expression of cathelicidin genes, we next sought to identify transcription factors driving such patterns. We analyzed the regulatory regions of *Camp3* and *Camp4*, two neighboring genes sharing regulatory elements and co-expressed at high levels in the same cells (Figs. 2A and C, and fig. S5). First, we carried out motif enrichment analysis on putative CREs and then cross-referenced the results with our gene expression data, as well as with publicly available ChIP data for human and mouse (figs. S7 and S8, and data table S3). Through this filtering strategy, we identified 5 transcription factors (*Spi1*, *Fli1*, *Runx1*, *C/ebpδ*, and *C/ebpε*) which were robustly expressed in P0 livers (fig. S7). To test whether these transcription factors could bind to putative CREs, we performed a set of luciferase assays using our immortalized line of sugar glider dermal fibroblasts (*47*). Specifically, we co-transfected each of the five transcription factors with luciferase reporter vectors containing the putative *Camp3* CRE, which was used as a representative sequence. Quantification of fluorescence after 48 h revealed that, out of the different transcription factors tested, *C/ebpδ*, and *C/ebpε* drove significantly higher levels of luciferase activity, compared with the GFP control vector (fig. S7 and table S4). Indeed, all highly expressed cathelicidins -*Camp3*, *Camp4*, *Camp6*, and *Ngp* contained a *C/ebp* transcription factor binding motif in their promoter or in a nearby intergenic peak (Fig. 2B). Notably, however, the putative shared CRE found to interact with both *Camp6* and *Ngp* in the RCMC analysis described above (Fig 2D), did not contain a *C/ebp* motif (Fig. 2B). Therefore, additional transcription factors may be involved in facilitating local enhancer-promoter interactions regulating *Camp6* and *Ngp*.

Taken together, our open chromatin data coupled to our RCMC data indicate that cathelicidin co-expression can take place via enhancer sharing. Moreover, interactions between regulatory sequences and different cathelicidin genes can take place both within and between Clusters. Lastly, our luciferase data suggest that, while *C/ebp* transcription factors are key regulators of cathelicidin expression, the presence of additional transcription factors is likely required to coordinate the regulatory control of members of the gene family.

### The function of sugar glider cathelicidins

The large repertoire of cathelicidin genes observed in marsupials, compared to mice and humans, raises the intriguing possibility that this gene family has been fine-tuned by natural selection such that different genes have acquired distinct roles. To test this, we next performed a variety of *in vitro* and *in vivo* assays aimed at characterizing the function of sugar glider cathelicidins. First, we investigated whether cathelicidins, like their eutherian homologues, exhibit antimicrobial properties, and the extent to which different genes vary in this ability. *In vivo*, cathelicidins are cleaved into mature peptides by serine proteases (*49*, *50*). Therefore, we predicted the most plausible cleavage sites and then chemically synthesized the mature peptides for 7 sugar glider cathelicidins (CAMP1, CAMP3, CAMP4, CAMP7, CAMP8, CAMP9, and CAMP10) and for NGP (Fig. 3A and table S2) (*34*, *51*, *52*). We did not synthesize mature peptides of CAMP6, due to its considerable length (137 AA), nor of the putative pseudogenes *Camp2* and *Camp5*. Most of the synthesized peptides were predicted to have a helical secondary structure, like other canonical cathelicidin peptides (fig. S9).

**Figure 3.**
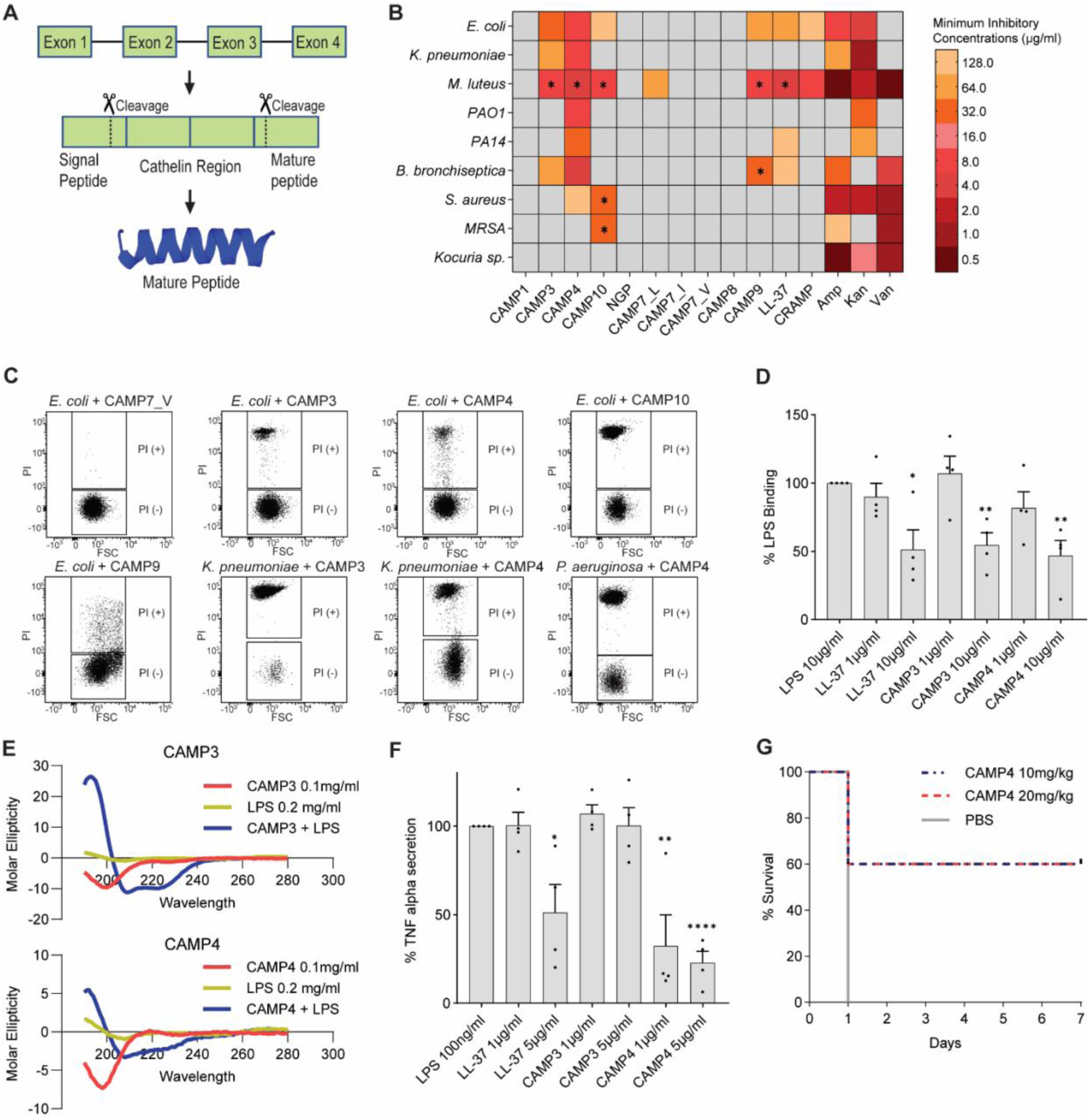
Function of sugar glider cathelicidins. (**A**) Schematic representation of cathelicidin peptide processing. Cathelicidins, composed of a signal peptide, a cathelin region, and a mature peptide, get cleaved into mature peptides by proteolytic proteins. (**B**) Heat map showing the minimal inhibitory concentrations (MICs) of CAMPs against different types of bacteria. Asterisks denote the existence of a higher peptide concentration where percent inhibition was lower than 90%. (**C**) Flow cytometry analysis of propidium iodide incorporation. Upon bacterial membrane disruption, PI intercalates with DNA, emitting a fluorescence signal. X-axis and Y-axis represent forward scatter and fluorescence intensity of propidium iodide, respectively. (**D**) Bar graph representing the fluorescence intensity, as measured via flow cytometry, of macrophages co-incubated with LPS-FITC and different peptides. Y-axis represents the percent of the median fluorescence intensity relative to the PBS control. Statistical significance was assessed using an unpaired *t*-test (n=4; *: p<0.05, **: p<0.01, ***:p<0.001, ****:p<0.0001). (**E**) Graphs showing the shift in the circular dichroism spectra of CAMP3 and CAMP4 upon incubation with LPS. The existence of negative bands around 220nm and the positive band at 193nm indicate the formation of an alpha-helix caused by the interaction with LPS. (**F**) Bar graph showing the amount of TNF-alpha secreted from LPS-stimulated macrophages upon co-incubation with peptides. Values are displayed relative to the PBS control. Statistical significance was assessed using an unpaired *t*-test (n=4; *: p<0.05, **: p<0.01, ****:p<0.0001). **(G)** Result from the *in vivo* mouse survival assay. Mice infected with *E. coli* were treated with CAMP4 or PBS control (n=5).

We then performed broth micro-dilution antimicrobial assays against *Escherichia coli, Klebsiella pneumoniae, Micrococcus luteus, Pseudomonas aeruginosa, Bordetella bronchiseptica,* and *Staphylococcus aureus*, common mammalian pathogens that were previously found in the pouch of wild marsupials (*15*, *16*), as well as against *Kocuria sp*., which was isolated from the pouch and skin swabs of female sugar gliders within our colony (Fig. 3B and table S5). For comparison, we included synthesized peptides corresponding to the human (LL-37) and mouse cathelicidins (CRAMP), as well as the antibiotics ampicillin, kanamycin, and vancomycin, all of which are known to have strong antimicrobial activity (*53*, *54*). We found that CAMP3, CAMP4, CAMP9, and CAMP10 exhibited strong antimicrobial activities against various bacteria, while CAMP1, CAMP7, CAMP8 and NGPs had low or no antimicrobial activities (Fig. 3B and table S5). Among the peptides with strong antimicrobial activity, CAMP4 was the strongest, showing marked efficacy against *E. coli*, *K. pneumoniae*, *M. luteus*, *B. bronchiseptica* and *P. aeruginosa* with minimum inhibitory concentrations (MICs) as low as 4 µg/ml, which are lower or comparable to the values of both human and mouse cathelicidins against these microbes (Fig. 3B and table S5). Additionally, the MIC of CAMP4 against particular bacterial species was comparable to that of common antibiotics (e.g., ampicillin against *E. coli*) (Fig. 3B and table S5). Importantly, the MIC ranges for the strongest cathelicidins (i.e., CAMP3, CAMP4, and CAMP10) were only minimally cytotoxic to mammalian cells, with cell viabilities higher than 90% for both immortalized sugar glider fibroblasts and J774.1 murine macrophages (fig. S10). Notably, none of the cathelicidins showed any antimicrobial activity against the sugar glider isolate *Kocuria sp* (Fig. 3B and table S5), suggesting that these potentially symbiotic microbes are resistant to the host’s AMPs. These results indicate that multiple sugar glider cathelicidins exhibit strong antibacterial properties against common mammalian pathogens, albeit to a varying extent depending on the cathelicidin and bacterial species being tested.

A common mechanism of action for mammalian AMPs is through disruption of bacterial membranes (*55*, *56*). To test whether this mechanism mediates antimicrobial activity of sugar glider cathelicidins, we performed a propidium iodide absorption assay (*57*). If bacterial membranes are disrupted by an antibacterial agent, propidium iodide will penetrate and intercalate with DNA, emitting a fluorescent signal that can be measured via flow cytometry (Fig. 3C). We applied this assay to *E. coli*, *K. pneumoniae*, and *P. aeruginosa* PAO1, using cathelicidin peptides that showed antimicrobial activities against each of the bacterial species in the broth microdilution assay (*E. coli*: CAMP3, CAMP4, CAMP9, and CAMP10; *K. pneumoniae*: CAMP3 and CAMP4; *P. aeruginosa* PAO1: CAMP4), along with the non-antimicrobial cathelicidin, CAMP7, as a negative control (Fig. 3C). We found that all four antimicrobial cathelicidins were able to disrupt bacterial membranes, while CAMP7 was not (Fig. 3C).

In addition to direct microbicidal activity, human and mouse cathelicidins (i.e., LL-37, CRAMP) have immunomodulatory properties, including chemotaxis, anti- and pro-inflammatory effects, and the promotion of Th17 differentiation (*58–63*). To establish whether sugar glider cathelicidins were capable of immunomodulation, we first incubated murine macrophages (J774.1) with FITC-conjugated LPS in the presence of cathelicidins or a vehicle control, and measured the resulting fluorescence intensity using flow cytometry. Among the peptides tested, CAMP3 and CAMP4 significantly reduced the binding of LPS to macrophages (Fig 3D and fig. S11). This effect was caused by direct binding of CAMP3 and CAMP4 to LPS, as determined by circular dichroism analyses (Fig 3E) (*64*). Next, we used ELISA to measure the amount of tumor necrosis factor alpha (TNF-alpha, a pro-inflammatory cytokine) released by murine macrophages upon LPS incubation, in the presence of LPS-binding cathelicidins or a vehicle control. Our results showed that CAMP4 significantly reduced TNF-alpha secretion compared to the vehicle control (Fig. 3F).

Finally, we sought to evaluate the *in vivo* efficacy of cathelicidins in a murine sepsis model (*65*, *66*). To this end, we tested whether CAMP4, the peptide that showed the strongest response in our antimicrobial assays as well as potent immunomodulatory activity, enhanced the survival of mice infected with *E. coli.* We intraperitoneally injected 5×10^7^ CFUs of *E. coli* into laboratory mice and treated them with one of two different concentrations of CAMP4 or a vehicle control (PBS). Strikingly, we found that while all mice treated with PBS died within 7 days, mice treated with both 10mg/kg and 20mg/kg of *Camp4* showed 60% survival rate (3 out of 5 mice survived) (Fig. 3G). This demonstrated that *Camp4* is effective in protecting mice from lethal sepsis, further corroborating our *in vitro* results.

### Evolution of sugar glider cathelicidins

The immunological significance of cathelicidins and the large number of these genes in marsupial genomes (*22*, *37*, *40*) prompted us to investigate the evolutionary history of this gene family. First, we examined patterns of divergence among sugar glider cathelicidins. We conducted pairwise comparisons of amino acid sequences among all sugar glider cathelicidins encoding functional proteins and found that these genes have experienced marked sequence diversification in the 4^th^ exon, which is the region encoding the mature effector peptide (Fig. 4A). Moreover, comparisons among *Camp 3*, *Camp4*, and *Camp10*, the three most similar genes, indicated that the nucleotide sequence identity of the 4^th^ exon (37.8%) was significantly lower than those of 3 introns (85.5%, 91.7%, and 84.7%, respectively) (Fig. 4B).

**Figure 4.**
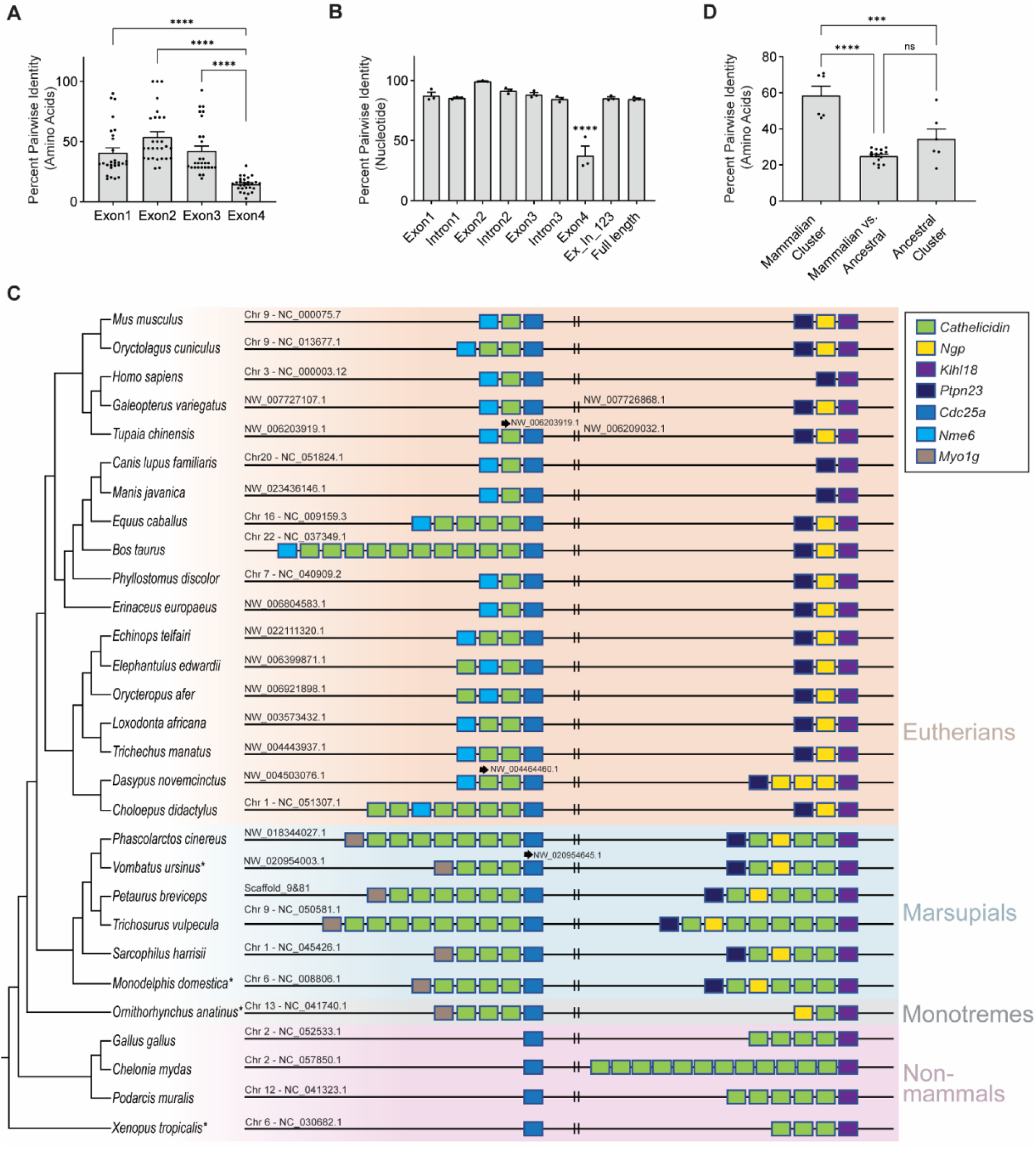
Evolution of the cathelicidin gene family. (**A**) Percent sequence identity (amino acids) among the exons of eight sugar glider cathelicidins. Statistical significance was assessed using an one-way ANOVA with Tukey’s multiple comparisons (****p<0.0001). **(B)** Percent sequence identity (nucleotide) among introns and exons of *Camp3*, *Camp4*, and *Camp10.* Statistical significance was assessed using a one-way ANOVA and post-hoc pairwise comparisons were carried out using a Bonferroni correction; *: p<0.05, ****p<0.0001). (**C**) Synteny map of cathelicidin gene clusters across mammals and non-mammalian vertebrates. Asterisks denote species in which cathelicidins were present outside the shown region. **(D)** Percent Sequence identity (amino acids) within and between cathelicidin clusters. Statistical significance was assessed using a one-way ANOVA and post-hoc pairwise comparisons were carried out using a Bonferroni correction (p***<0.001, p****<0.0001).

Motivated by the extensive number of genes observed among marsupial cathelicidins, we expanded our evolutionary analysis to search for broader evolutionary patterns among tetrapods. To achieve this goal, we annotated cathelicidins and *Ngp* genes in publicly available genomes of 24 additional mammals (table S6). In addition, we included cathelicidin sequences of four non-mammalian tetrapods as outgroups (table S6). Then, we aligned sequences along with syntenic marker genes for cluster identification (Fig. 4C). Our analysis revealed that marsupials and monotremes are the only tetrapods with two clusters of cathelicidins, while eutherian mammals and non-mammalian tetrapods each have a single cluster (Fig. 4C). Notably, the cathelicidin cluster present in eutherian mammals is distinct from the cluster found in non-mammalian tetrapods as indicated by their syntenic relationships with surrounding genes and the phylogenetic relationships of their constituent cathelicidins. This observation suggests that a duplication event of the cathelicidin gene cluster took place in the mammalian ancestor, prior to the divergence between monotremes and therian mammals. Marsupials and monotremes have retained both the ancestral tetrapod and novel mammalian clusters, while eutherian mammals lost the ‘ancestral’ cluster (Cluster 2 in sugar gliders) shared with non-mammalian tetrapods, retaining only the ‘mammalian’ cluster (Cluster 1 in sugar gliders). Further pairwise sequence comparisons among sugar glider cathelicidins revealed that genes within the ‘mammalian’ cluster show higher identity than those within the ‘ancestral’ cluster, reinforcing that the mammalian cluster has likely arisen more recently (Fig. 4D).

Lastly, our comparative annotation and synteny map suggests that *Ngp* is a gene that exists only in mammals. We carried out a phylogenetic analysis of 151 tetrapod cathelicidin and 28 *Ngp* amino acid sequences and found that all *Ngp* sequences formed a monophyletic group (fig. S12 and data table S4). Furthermore, when we blasted amino acid sequences of representative mammalian *Ngp* genes against non-mammalian protein database, we found that the top hits were cathelicidins, indicating that *Ngp* evolved from a cathelicidin in early mammal or pre-mammalian synapsid diversification (table S7).

Overall, our phylogenetic characterization of cathelicidin and *Ngp* genes indicates that mammalian cathelicidins have experienced multiple evolutionary events, including both duplication and loss of gene clusters, as well as the derivation of a new gene, *Ngp*. The genomic expansion and the retention of two different cathelicidin clusters suggest that cathelicidins likely played a crucial role in shaping marsupial immune defenses throughout evolution.

## Discussion

At birth, the transition to a microbe-rich extrauterine environment presents mammalian neonates with significant challenges, due to their naïve immune systems. This is particularly true of the marsupials, which complete the development of lymphoid organs postnatally (*13*). The exceptionally altricial state of marsupial neonates increases their reliance on innate immunity during the first days and weeks of life. Here, we show that cathelicidins represent a key component of the early postnatal immunological defense system in marsupials. Through single-cell transcriptomic analysis of the P0 sugar glider liver, we have characterized the cellular composition and the repertoire of genes expressed in the neonatal marsupial hematopoietic tissue. We present evidence that several cathelicidin genes are highly co-expressed in neutrophils, and that this process is partly regulated by enhancer sharing and long-range physical interactions between loci found in the two clusters. Moreover, we show that sugar glider cathelicidins possess immunomodulatory properties, can directly kill bacteria by disrupting their cell walls, and are able to protect mice from lethal sepsis. Lastly, our comparative genomic analysis revealed that marsupials and monotremes—both of which give birth to highly altricial neonates (*67*)—retain both ancestral and mammalian cathelicidin clusters, while eutherians and non-mammalian tetrapods possess a single cluster.

By studying sugar gliders, a species suitable for captive breeding, and by combining transcriptomics, epigenomics, functional experiments, and comparative genomics, our study considerably advances our understanding of the regulation, function, and evolution of marsupial neonate immune protection. Nonetheless, there are several questions that remain to be addressed. First, although our functional studies used bacterial species previously identified in other marsupials and in our captive colony, it would be crucial to also characterize the pathogens found in the sugar glider’s native environment. This would allow us to test whether sugar glider cathelicidins are effective against the pathogens they naturally encounter. Second, while many sugar glider cathelicidins possess immunomodulatory and antimicrobial functions, others, such as *Camp6*, *Camp7*, and *Ngp*, do not, despite being highly expressed. As such, functionally characterizing the genes that make up Cluster 2 may yield fascinating new insights. For example, due to its unique expression pattern, *Camp7* is likely to have non-canonical functions different from those of other cathelicidin genes. Third, while our assays demonstrate that sugar glider cathelicidins are functionally important, the extent to which these genes protect sugar glider neonates remains unknown. Genome editing approaches in sugar gliders will eventually allow us to systematically study the function of each cathelicidin gene through loss-of-function experiments (*68*).

In addition to motivating these questions, our results pave the path towards studying foundational concepts in mammalian gene regulation, ecology, and evolution. For example, the high-resolution regulatory interactions between the two cathelicidin gene clusters provide a framework for identifying genomic elements, transcription factors, and long-range co-expression determinants between distant genomic loci. Furthermore, marsupial cathelicidin evolution exemplifies how specific gene duplication events reflect key life history aspects of a taxon. As different mammalian species exhibit a broad spectrum of developmental maturity at birth (*67*), comparative genomic analysis across a range of altricial and precocial taxa will further illuminate the relationship between life history and the evolution of immune-related gene families.

## Materials and Methods

### Single cell RNA sequencing (scRNA-Seq)

A liver of a postnatal day 0 (P0) sugar glider neonate was digested with 0.02% collagenase II for 30 minutes at 37°C with gentle rotation. Primary cells were filtered with a 70um cell strainer and then centrifuged for 4 minutes at 800g at 4°C. The resulting cell pellet was suspended in 1 x PBS containing 0.04% FBS. Cell viability and number were assessed with a hemocytometer using trypan blue. A scRNA-seq library was prepared using the Chromium Single Cell 3-prime Library Prep Next GEM (v3.1) and was sequenced on an Illumina NovaSeq 6000 (28 × 90bp, paired-end), generating 508M reads. The resulting reads were mapped against the sugar glider genome using CellRanger-7.0.0. Cells containing more than 200 genes, but less than 10% of total mitochondrial genes, were kept for downstream analysis. UMI count data of surviving cells were normalized using Seurat v4.3.0 with SCTransform (*69*). We performed principal component analysis (PCA) and ran uniform manifold approximation and projection (UMAP) dimensional reduction technique (n=30 dimensions) using Seurat’s default parameters. Neighbors and clusters were identified with 30 dimensions and 1.0 resolution, respectively.

Clusters were annotated based on the following marker genes: Red blood cell lineages: *Hemgn*, *Klf1*, *Kit*, *Band3*, *Gpa*, and hemoglobin subunits (*70*, *71*); Hematopoietic stem cell clusters: *Cd34* and *Znf521* (*72*, *73*); Neutrophil lineage cells: *S100A8*, *Gfi1*, *Mpo*, *Prtn3*, *Elane*, *Ltf*, *Ngp*, *Csf3r*, *Prex1*, and *Pde4b* (*74–76*); Lymphoid cells: lymphoid marker genes *Cd3* subunits, *Blk*, and *Jchain* (*77*, *78*); Monocyte-macrophage lineages: *Maf*, *Mafb*, and *Cd80* (*79–83*); Megakaryocytes: *Tuba8* and *Gp1ba* (*84*, *85*); Mast cells: *LOC110197119* (mas-related G-protein coupled receptor member H-like) and *Ms4a2* (*86*, *87*); and Eosinophils: *Epx* and *Il5ra* (*88*, *89*).

Differential expression analysis was conducted using the FindMarkers() function of the Seurat package. Normalized gene expression values were visualized using FeaturePlot(). Co-expression plots were visualized using FeatureScatter().

### Annotation of cathelicidins and *Ngp*

After compiling sequences of 79 mammalian cathelicidin and cathelicidin-like sequences from NCBI, we blasted protein sequences of all cathelicidins against the previously constructed *de novo* sugar glider transcriptome (*44*). For targeted gene annotation, we next blasted both 79 mammalian cathelicidin protein sequences and *de novo* sugar glider transcript hits with e-values lower than 10^-5^ against the sugar glider genome, and extracted sequences 1 mb upstream and downstream of the resulting blast hits.

Using *de novo* transcript hits, as well as mRNA and protein sequences of 79 mammalian cathelicidins as EST evidence, we ran Maker3 (*90*) on extracted genome sequences (six rounds) using the Augustus human gene prediction model with snap hmm building (*91*, *92*). To annotate cathelicidin genes and *Ngp* in 24 additional mammalian species (table S6, *Ornithorhynchus anatinus*, *Monodelphis domestica*, *Sarcophilus harrisii*, *Trichosurus vulpecula*, *Vombatus ursinus*, *Phascolarctos cinereus*, *Choloepus didactylus*, *Dasypus novemcinctus*, *Trichechus manatus latirostris*, *Loxodonta africana*, *Orycteropus afer afer*, *Elephantulus edwardii*, *Echinops telfairi*, *Erinaceus europaeus*, *Phyllostomus discolor*, *Bos taurus*, *Equus caballus*, *Manis javanica*, *Canis lupus familiaris*, *Tupaia chinensis*, *Galeopterus variegatus*, *Homo sapiens*, *Oryctolagus cuniculus cuniculus, Mus musculus*), we used mammalian cathelicidin and NGP sequences as EST evidence and performed the same targeted gene annotation using a single round of the Maker3 run. Resulting annotations were manually compared with NCBI reference annotations to accurately count the number of genes. For 4 non-mammalian tetrapods (*Xenopus tropicalis, Podarcis muralis, Chelonia mydas, Gallus gallus*), we primarily relied on the NCBI and ensemble annotations, since our Maker3 annotation based on mammalian cathelicidins did not provide accurate annotations. Specifically, focusing on the ancestral cluster locus, we looked for genes annotated as cathelicidins or cathelicidin-like. Then, to find additional cathelicidins, we blasted protein sequences of these genes against the protein database of the respective species.

For further validation of sugar glider cathelicidin and *Ngp* sequences, we generated cDNA from RNA extracted from lung, liver, stomach, kidney, intestine, and brain of a ∼P20 (1.35g) joey. PCR amplifications followed by Sanger sequencing confirmed the sequences of six cathelicidins and of *Ngp*.

To infer the genomic location of *Camp10*, we first acquired protein sequences of cathelicdiins from the leadbeater’s possum (*Gymnobelideus leadbeateri*, Genome assembly LBP_v1:GCA_011680675.1), a sister species of the sugar glider, by lifting-over gene model from sugar glider cathelicidin annotation (*47*, *93*). We then conducted reciprocal protein blast of cathelicidins from the leadbeater’s possum and the sugar glider, as well as a nucleotide blast search of a 7000bp region of scaffold 81, which contains cathelicidin 10, against the genome of the leadbeater’s possum (fig. S3).

### Phylogenetic tree construction

Sugar glider cathelicidin and *Ngp* nucleotide sequences aligned with MAFFT v7.475 were fed into RAxML for the construction of unrooted phylogenetic trees with 1000 bootstrap values and GTRGAMMA model (*94*, *95*). To build a comprehensive phylogeny of cathelicidins and *Ngp*, we compiled 151 cathelicidins and 28 NGP sequences from NCBI (data table S4). We built a phylogenetic tree with MEGA-X using cathelicidin and NGP sequences aligned with the built-in muscle aligner (100 bootstraps and JTT model)(*96*). All trees were visualized with FigTree (*97*).

### Bulk RNA sequencing

For bulk RNA sequencing, sugar glider liver samples were collected from P0 neonates (n = 3), ∼P10 joeys (0.45g, 0.55g, 0.56; n = 3), and adults (n = 3). Additionally, we collected bone marrow samples from adult tibia and femur (n = 3), as previously described (*98*). Briefly, hind limb bones were cut (around the kneecap) and placed into 0.5 mL tubes. Using a sterile pin, a hole was poked in the bottom of the tube and it was placed inside a 1.5 mL container with 100 µL of PBS. The bone marrow sample was then collected by centrifugation (8000g, 30s at RT). The RNeasy mini kit (QIAGEN) was used for RNA extraction following the supplier’s protocol. We used 250ng of RNA to prepare libraries with NEBNext Ultra II Directional RNA Library Prep kit. Libraries were sequenced to a median depth of 27M reads on the NovaSeq 6000 (2 × 65bp). Demultiplexed reads were filtered and trimmed using Trimmomatic 0.39 with parameters: MINLEN:25 AVGQUAL:20 (*99*). Resulting reads were mapped against the sugar glider genome using STAR-2.7.8a (*100*). RPKM (Read per kilobase per million reads) values were calculated based on read counts acquired from featureCounts (*101*). RPKM values were visualized using GraphPad Prism.

### ATAC sequencing

Single cell suspensions were generated from a liver of a P10 joey as described for scRNA-seq. For each replicate (n = 3), library preparation was done using 100,000 cells, following the Omni-ATAC protocol (*46*). Briefly, following lysis on ice for 3 minutes, cells were incubated with TDE1 transposase (Illumina) for 1h at 37°C and then purified with the Zymo DNA Clean and Concentrator-5 kit (Zymo). Illumina sequencing adaptors and barcodes were added to the DNA fragments. The resulting libraries were sequenced on NovaSeq SP flowcell as 61 nt reads, generating 59M, 66M, 64M reads for each of the 3 libraries. Raw ATAC reads were trimmed by NGmerge and then mapped to the genome using Bowtie2 (*102*). Peaks were called using MACS2 with following parameters: --nomodel -q 0.05 --keep-dup all --shift −100 --extsize 200 -g 2456432000 –nolambda (*103*). IDR (Irreproducible Discovery Rate) was used to assess peak calls concordance (*104*). Only peaks called in at least two out of three pairwise IDR analyses were considered for downstream analyses. Peak data were visualized in IGV (*105*).

### RCMC library construction

Bone marrow samples were collected from adult sugar gliders (n =2) as described above. Cells were incubated with 0.2% NaCl for 30 seconds and red blood cells were removed by incubating the sample with 1.6% NaCl for 30 seconds. Micro-C libraries were generated from 3.2M and 3.3M cells, using previously described methods (*48*, *106–110*). Briefly, samples were crosslinked with 4% formaldehyde (20 minutes and 500 rpm at room temperature), followed by the secondary crosslinking in 1mL of freshly prepared 3mM DSG (ThermoFisher 20593) and ESG (ThermoFisher 21565) in PBST (40 minutes, 500 rpm at room temperature). The reaction was quenched by adding 250 μL 2M Tris-HCl pH7.5 for 5 minutes, washed twice with PBS containing 3% BSA, snap-frozen, and stored at −80℃. For library construction, samples were digested with micrococcal nuclease (concentration determined by titration) (20 minutes, 1000 rpm at room temperature) in MB1 buffer (50mM NaCl, 10mM Tris-HCl pH7.5, 5mM MgCl2, 1mM CaCl2, 0.2% NP-40, and Protease Inhibitor Cocktail (Sigma 11697498001), followed by heat inactivation with 5mM EGTA at 65C℃ for 15 minutes. Samples were washed 3 times with MB2 buffer (50mM NaCl, 10mM Tris-HCl pH7.5, 10mM MgCl2 and 1mM CaCl2), and then treated with 0.5U/μL T4 PNK (New England Biolabs M0201) (30 minutes, 1000 rpm at 37 ℃) in End-prep buffer (1X NEB buffer 2.1 (New England Biolabs B7202), 2.5mM ATP, and 5mM DTT). After T4 PNK treatment, samples were incubated with Klenow Fragment (New England Biolabs M0210) at a final concentration of 0.5U/μL (15 minutes, 1000 rpm at 37 ℃). End-labeling cocktails were added to the samples to make the final concentration 50μM Biotin-dATP (Jena Bioscience NU-835-BIO14), 50μM Biotin-dCTP (Jena Bioscience #NU-809-BIOX), 50μM dGTP, 50μM dTTP, 0.3X T4 DNA ligase buffer (New England Biolabs B0202), and 80μg/mL BSA. Samples were incubated (45 minutes, 1000 rpm at room temperature) and heat inactivated with 5mM EDTA (15 minutes at 65 ℃). Samples were washed 3 times with MB3 buffer (40mM Tris-HCl pH7.5 and 10mM MgCl2) and then treated with 20U/μL T4 DNA ligase (New England Biolabs M0202) (16 hrs, 400 rpm at 16 ℃) in ligation buffer (1X T4 DNA ligase buffer (New England Biolabs B0202) and 200μg/mL BSA. After ligation, samples were treated with 4U/μL Exonuclease III (New England Biolabs M0206) in 1X NEB buffer 1 (New England Biolabs B7001) (30 minutes, 1000 rpm at 37 ℃), followed by addition of 0.5 mg/mL RNAseA (Thermal Fisher EN0531) (30 minutes, 1000 rpm at 65 ℃). Samples were then treated with 0.5mg/mL ProteaseK in 1% SDS (overnight at 65℃). After protease K treatment, DNA libraries were extracted and purified from samples using standard Phenol:Chloroform extraction and Ethanol precipitation. Libraries were sheared to 200bp fragments using Covaris ME220 (Duration 130s, Peak Power 70W, Duty Factor 20%, Cycles per Burst 1000). Libraries were then bound to Dynabeads™ MyOne™ Streptavidin C1 beads (Thermal Fisher 65001) and washed following the manufacturer’s instructions. Libraries were end repaired and ligated to adaptors using NEBNext Ultra II library prep kit (New England Biolabs E7645). Next, libraries were amplified with NEBNext® Multiplex Oligos for Illumina® (Dual Index Primers Set 2) (New England Biolabs E7780) and KAPA HiFi HotStart ReadyMix (KAPA Biosystems KK2601), and then purified with Ampure XP beads (Thermal Fisher #NC9959336). The Micro-C libraries were then subject to quality control with Qubit and bioanalyzer. For RCMC, the custom probe sets were designed using the sequence of the target capture region (HiC_scaffold_9: 107950000-109750000; HiC_scaffold_81: 114775). Micro-C libraries were hybridized to the probe sets, pulled down, and amplified using Twist Standard Hyb and Wash Kit v2 (Twist Biosciences 105560). The final RCMC libraries were cleaned up with Ampure XP beads (Thermal Fisher #NC9959336) and subjected to paired-end sequencing on an Illumina Novaseq S1 100 nt Flowcell, yielding a total of 446M reads.

### RCMC data processing

Micro-C data were aligned to custom genomes with BWA-MEM using parameters -S -P −5 -M (*111*). The resulting BAM files were parsed, sorted, de-duplicated, filtered, and split with Pairtools (https://github.com/open2c/pairtools). We removed pairs where only one half of the pair could be mapped, or where the MAPQ score was less than three. The resulting files were indexed with Pairix (pairtools function). The files from replicates were merged with Pairtools (https://github.com/open2c/pairtools) before generating 100bp contact matrices using Cooler (*112*). Finally, balancing and mcool file generation was performed with Cooler’s Zoomify tool, and visualized on Higlass (*113*).

### Transcription factor motif analysis

FIMO (Find Individual Motif Occurences) motif scan was first conducted against putative enhancers of *Camp3* and *Camp4* as the expression levels, sequence identity of both genes and enhancers are similar to each other, suggesting the existence of shared regulatory mechanism (*114*). Featureplots and violinplots from scRNA-seq datasets as well as RPKM values from RNA-seq datasets were cross-referenced for narrowing down candidates. *C/ebpδ*, *C/ebpε*, *Spi1*, *Fli1*, *Runx1*, *Foxp2*, and *Nfatc3* stood out as initial candidates but *Foxp2* and *Nfatc3* were excluded as they did not bind to human and mouse cathelicidin locus (fig. S8) (*115–117*). Further FIMO motif scans were conducted on additional putative regulatory peaks to search for *C/ebp* family binding sites.

### Luciferase assays

Sequences of a putative enhancer region of *Camp3* was PCR amplified from sugar glider genomic DNA (table S1) and cloned into the pGL4.23 Luciferase Enhancer Reporter Vector using In-fusion cloning (Takara). Sequences for the different transcription factors and *Camp3* promoter were obtained either commercially (*Camp3 promoter*, *Spi1*, *Fli1*, *Runx1*, *Ikzf1* and *C/ebpε*; Twist Biosciences) or by extracting RNA from the bone marrow of an adult sugar glider and reverse transcribing it using qScript cDNA SuperMix (Quantabio) (*C/ebpδ*) (table S1). Transcription factor fragments were then cloned into the pCMV-GFP vector. A total of 2,000 immortalized sugar glider cells were seeded in a white 96 well plate (PerkinElmer) and transfected with the experimental constructs (200ng) as well as a control pGL4.74 Renilla Reporter Vector (20ng) using 0.3µl of Lipofectamine (Invitrogen). After 48 hours of incubation at 37°C, cells were harvested and analyzed with the DualGlo Luciferase Assay System (Promega). Luminescence was measured by Tecan Spark microplate reader. Experiments were performed in six replicates and statistically analyzed using an unpaired t-test in GraphPad Prism.

### Pouch swabbing

Sugar glider pouches were swabbed for 30 seconds with cotton swabs pre-moistened with ultra-pure water. Swabs resuspended in ultra-pure water were plated in LB media and incubated overnight at 37°C. White colonies that repeatedly grew from pouch swabs of multiple females were isolated and subject to DNA extraction (Zymo D6005). Bacterial species were identified by Sanger sequencing of the PCR amplified 16S rRNA gene (827F: 5’AGA GTT TGA TCC TGG CTC 3’, 1492R: 5’ TAC GGY TAC CTT GTT ACG ACT 3’).

### In vitro broth micro-dilution antimicrobial assay

The first Alanine or Valine found in the 4^th^ exon of CAMP1, CAMP3, CAMP4, NGP, CAMP7, CAMP9, and CAMP10, were predicted to represent the cleavage sites for the mature peptides (*34*). For CAMP8, due to the short length (15AA) and the lack of Alanine and Valine in the 4^th^ exon, we set the cleavage site to be the last Alanine in the 3^rd^ exon. For CAMP7, due to the short length of the predicted mature peptide after the Valine, we synthesized three versions of mature peptides with predicted cleavage sites located respectively in the first Leucine, the Isoleucine, and the Valine found in the 4^th^ exon. We did not synthesize mature peptides of CAMP6 due to its extensive length (137 AA), nor of the pseudogenes *Camp2* and *Camp5*. Secondary structures of synthesized mature peptides were predicted using PEP-FOLD3 (*118*).

We used broth micro-dilution assays (*119*) to measure minimal inhibitory concentration (MIC) of cathelicidins against *Escherichia coli* (ATCC 25933)*, Klebsiella pneumoniae* (ATCC 43816)*, Micrococcus luteus* (ATCC 4689)*, Pseudomonas aeruginosa* (PAO1, PA14)*, Bordetella bronchiseptica* (ATCC 10580)*, Staphylococcus aureus* (ATCC 29213, MRSA), and *Kocuria sp*. Briefly, we added 10^5^ CFU of bacteria from overnight cultures into each well of a 96-well plate (costar, Corning) with different concentrations of each peptide or antibiotics. 200µl of cation-adjusted Mueller-Hinton broth (CAMHB) was added to each well. Samples were incubated at 37°C and absorbance was measured at 0 h and 16 h using the Tecan Spark microplate reader (600 nm). For *M. luteus*, we measured absorbance at 43 hours as well. MIC was determined as the lowest concentration that showed no visible change after 16 hours (43 hours for *M. luteus*) incubation for all three replicates and showed > 90% mean percent inhibition based on absorbance measurements.

### Cell viability assay

Cell viability was measured using CellTiter-Glo 2.0 Assay (Promega) following the manufacturer’s protocol. Briefly, we incubated 10,000 cells in 100µl Dulbecco’s Modified Eagle Medium (DMEM) supplemented with 10% FBS, 1.5g/L sodium bicarbonate, 1mM sodium pyruvate, and 1% penicillin/streptomycin mix overnight (37°C) with different concentrations of cathelicidins and a SDS control. Subsequently, we added 100µl of CellTiter-Glo 2.0 reagent. After 2 minutes mixing and a 10 minute incubation at room temperature, we measured luminescence using Tecan Spark microplate reader. Experiments were done in triplicate.

### Propidium Iodide absorption assay

Bacteria were grown overnight to log phase. 10^6^ CFU of bacteria were incubated for 30 minutes at 37°C with different concentrations of cathelicidin peptides in 200µl PBS. After a 2 minute centrifugation at 8000 rpm, pellets were washed with 200 µl PBS and resuspended in 1ml PBS. Propidium Iodide (PI) was added for a final concentration of 1 µg/ml. An LSRII flow cytometer (BD Biosciences) was used to record 50,000 events of PI uptake. 0.25%-1% SDS was used as a control.

### *In vivo* mouse infection

6-8 week old female mice (C57BL/6J, The Jackson Laboratory) were intraperitoneally injected with 5 x 10^7^ CFU of *E. coli* (ATCC 25922) in 100 µl of PBS. 30 minutes after the bacterial injection, mice were treated with either CAMP4 peptide (10mg/ml or 20mg/ml) or PBS vehicle. Treated mice were checked daily for 7 days. All experiments performed were approved by the IACUC committee at Princeton University.

### LPS binding assay

Mouse macrophage cells (J774A.1; ATCC TIB-67) were grown in supplemented DMEM media in 7.5% CO2 at 37°C. 2.5 x 10^5^ cells were incubated with 10 µg/ml LPS-FITC and different concentrations of antimicrobial peptides in 500 ml PBS for 30 minutes. After washing the cells using PBS, median fluorescence intensity was determined using the LSRII flow cytometer (BD sciences). The percentage difference from negative control was statistically analyzed with unpaired-t tests in GraphPad Prism. Experiments were done in quadruplicate.

### Circular dichroism

The secondary structure of CAMP3 and CAMP4, which showed LPS-binding activity in at least one of the two tested concentrations, were determined using circular dichroism spectrometer (Chirascan, Photophysics). Measurements were taken in a rectangular cuvette, using an emission range of 180 to 280nm (in 1nm increments). 0.1 mg/ml of peptides and 0.2 mg/ml of LPS in sterile water were used for measurements.

### TNFa secretion assay

ELISA assays were performed according to manufacturer’s instructions (R&D Systems DuoSet). Briefly, a 96-well plate (Corning Costar) was coated with anti-TNFa and incubated overnight at room temperature. The plate was then washed with Wash Buffer, blocked for 1 hour with Reagent Diluent, washed again, and the liquid aspirated. J774A.1 macrophages were incubated with LPS, cathelicidin, vehicle, or a combination of these for 24 hours and 100 µl of supernatant from the incubating macrophages or cytokine standard, in sequentially diluted concentrations ranging from 2000-32 pg/mL, were applied to the wells and incubated for 2 hours at room temperature before being washed and aspirated. Biotinylated anti-TNFa, streptavidin-HRP, and substrate solution were applied in sequence. The plate was then incubated in the dark for 2 hours or 20 minutes and subsequently washed and aspirated. The HRP-substrate reaction proceeded for 20 minutes before the Stop Solution halted it. The plate was then read for optical density at 450 nm and 540 nm. The percentage difference from the LPS control was statistically analyzed with unpaired-t tests in GraphPad Prism. Experiments were done in quadruplicate.

### Pairwise sequence identity analysis

Amino acid sequences of 8 cathelicidin genes (*Camp1, Camp3, Camp4, Camp6, Camp7, Camp8, Camp9,* and *Camp10*) were aligned with MAFFT v7.475 in a pairwise manner. Resulting alignments were fed into Sequence Manipulation Suite for identity analysis (data table 34) (*120*). For statistical analysis, separate pairwise analysis was conducted for each exon. In this analysis, we used the truncated, Maker3-annotated version of *Camp6* due to the extensive length of the predicted protein compared to the rest of the cathelicidin genes. Nucleotide sequences of *Camp3, Camp4,* and *Camp10* were further aligned and compared for detailed analysis. Statistical analysis was conducted in GraphPad Prism using one-way ANOVA test. Post-hoc pairwise comparisons were carried out using a Bonferroni correction.

## Acknowledgments

We thank the members of the Mallarino laboratory and the Donia laboratory for discussions and assistance with animal husbandry; Princeton LAR (K. Gerhart, G. Barnett and J. McGuire, D, D. L. Aler, R. Franco, P. K. Cunningham, B. Tereszczyn) for help with animal husbandry and care; J. M. Miller and J. Arley Volmar from the LSI Genomics Core (Princeton) for help with library preparation, O. Perotti for help in the lab, and F. Rogers for help with statistics.

## Author Contributions

J.P., M.S.D., and R.M. conceived the project and designed experiments. J.P. and C.Y.F. conducted computational gene annotation, phylogenetic analyses, and tissue harvesting. W.K. conducted RCMC experiments. W.K., J.P., and Y.P. analyzed RCMC data. A.K. performed ELISA experiments and analyzed the resulting data. J.P. conducted all the other experiments and data analyses. J.P., M.S.D., and R.M. wrote the manuscript with the input from all authors.

## Data and materials availability

The scRNA-seq, RNA-seq, ATAC-seq, and RCMC data are submitted under NCBI BioProject accession code PRJNA1141454.

## Supplementary Materials

**Fig. S1.**
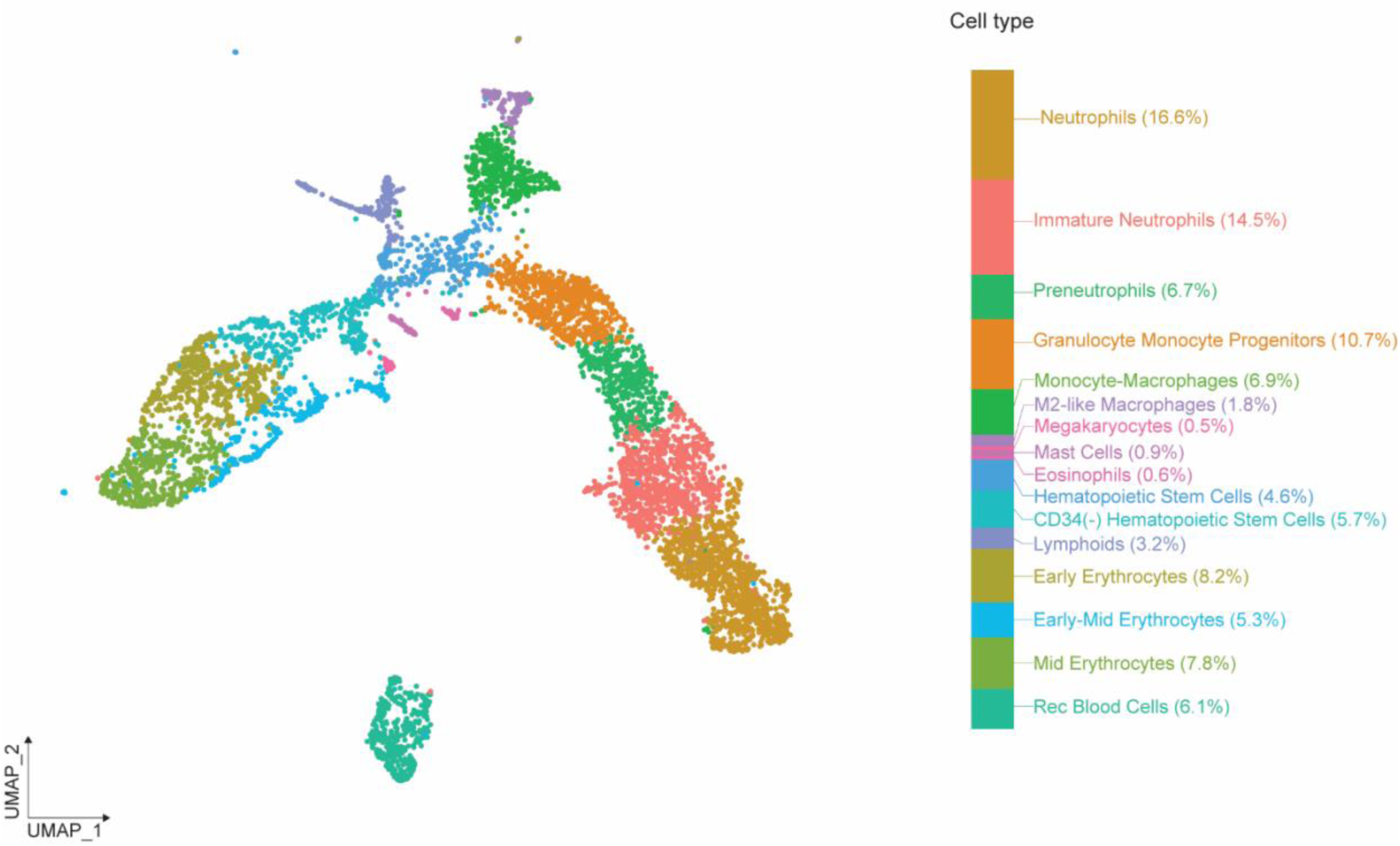
UMAP plot showing cell type clustering and composition of the sugar glider neonatal (P0) liver. Data was analyzed before the systematic annotation of cathelicidins.

**Fig. S2.**
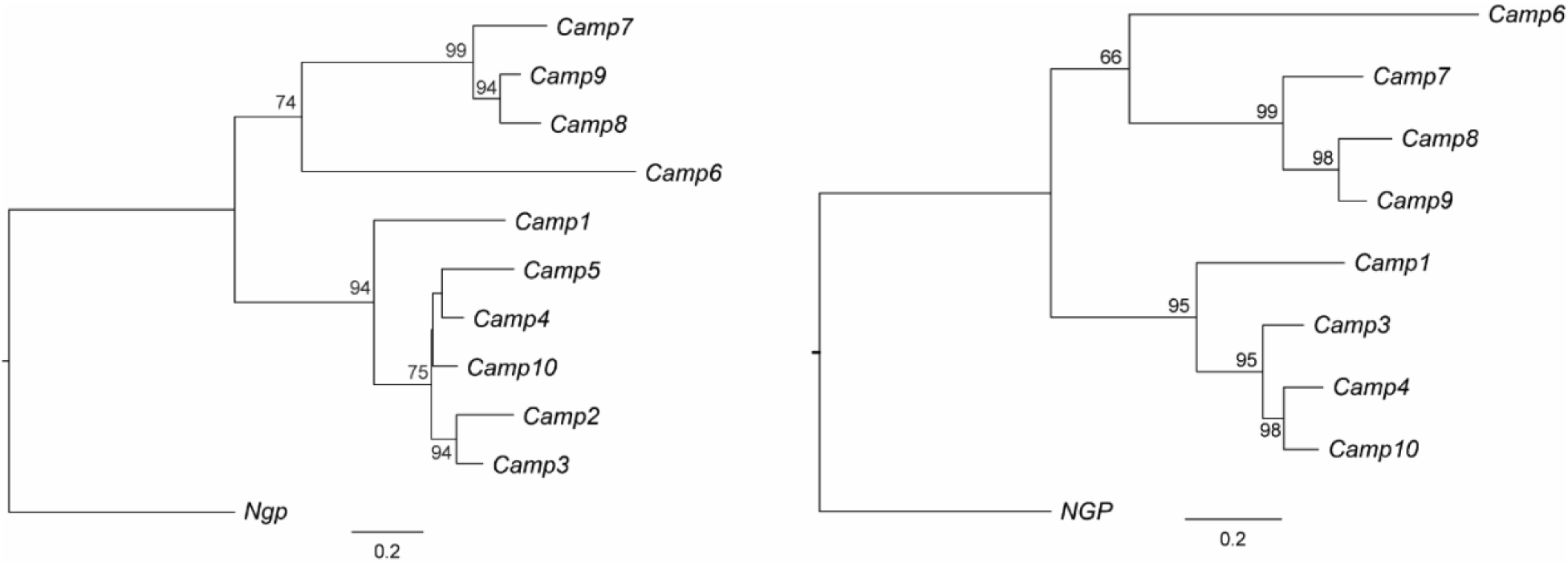
Evolutionary relationships among sugar glider cathelicidins. Maximum likelihood phylogenetic trees of cathelicidin genes with (left) and without (right) *Camp2* and *Camp5*. Only bootstrap values higher than 50 are shown.

**Fig. S3.**
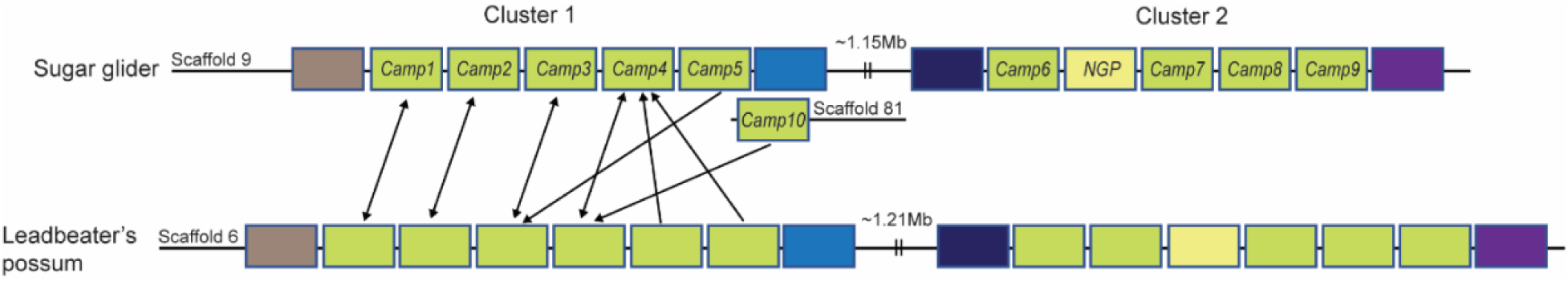
Schematics showing syntenic relationships between cathelicidins in sugar gliders and in the leadbeater’s possum. Reciprocal blasts were conducted to infer the location of *Camp10*. Arrows depict top blast hits.

**Fig. S4.**
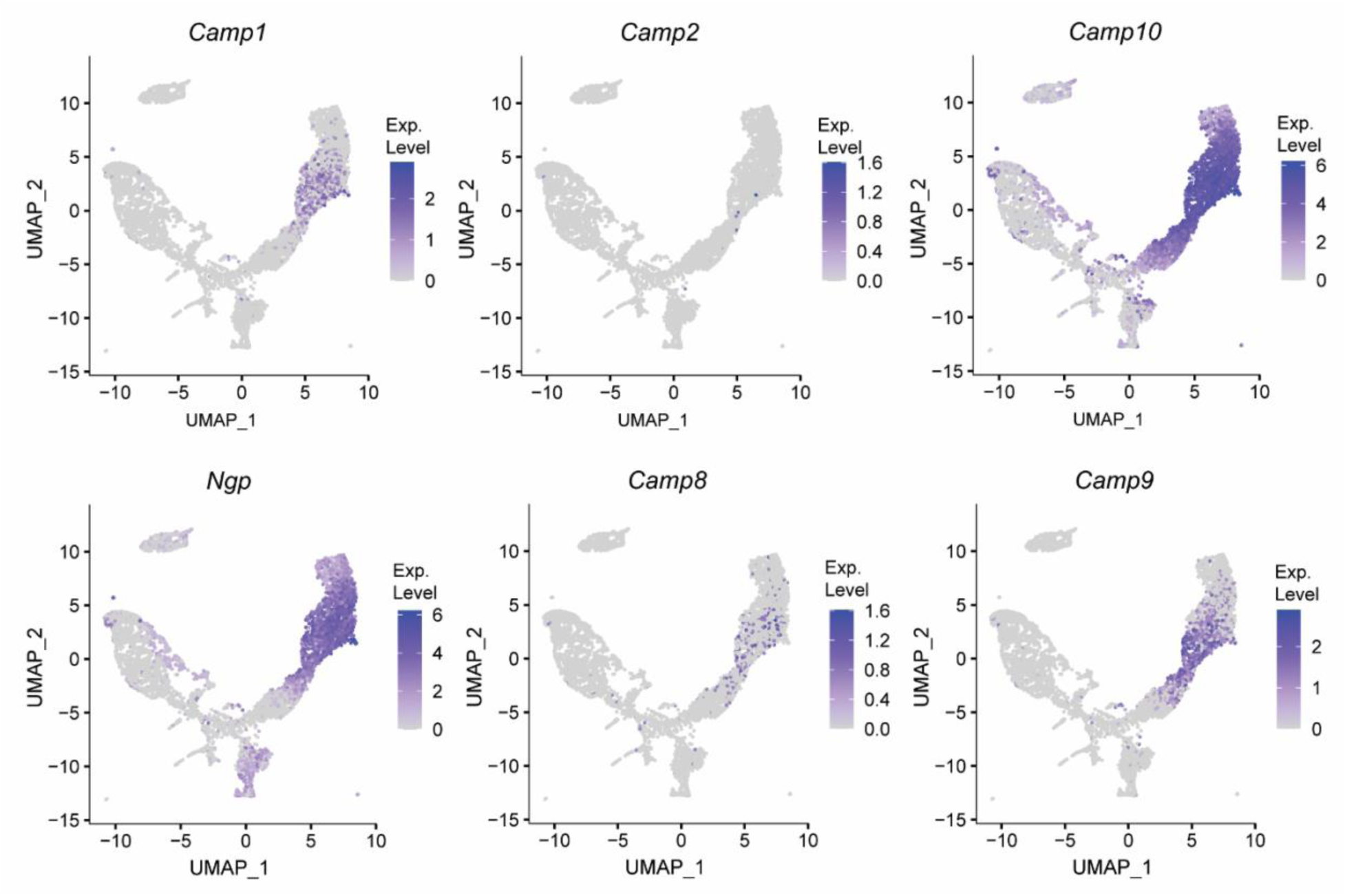
UMAP plots displaying the expression of selected cathelicidin genes.

**Fig. S5.**
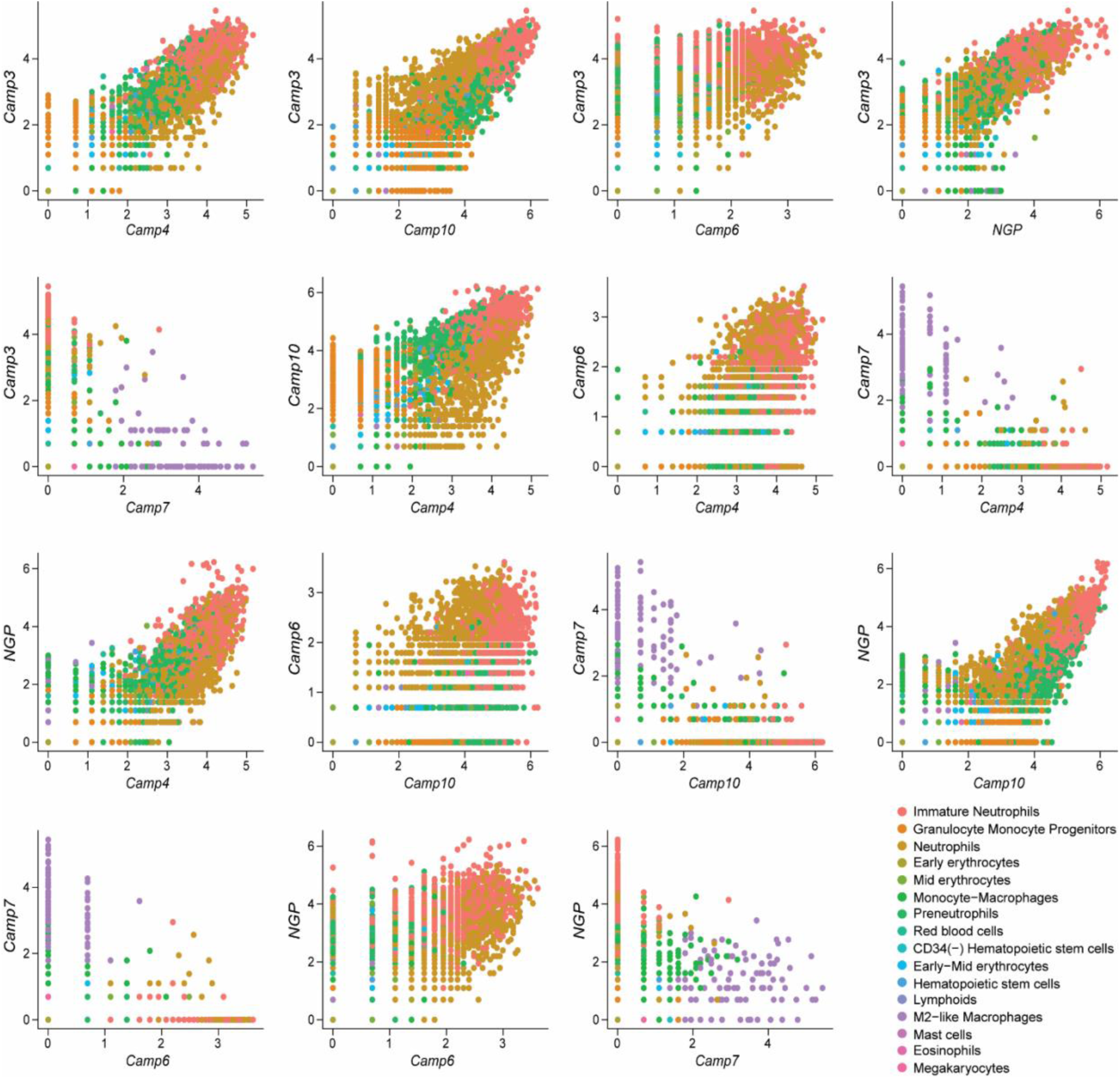
Co-expression plots of cathelicidin genes. Each dot represents a cell. *Camp3*, *Camp4*, *Camp10*, and *Camp6* were co-expressed in neutrophil lineage cells whereas Camp7 is not co-expressed with any other cathelicidin gene.

**Fig. S6.**
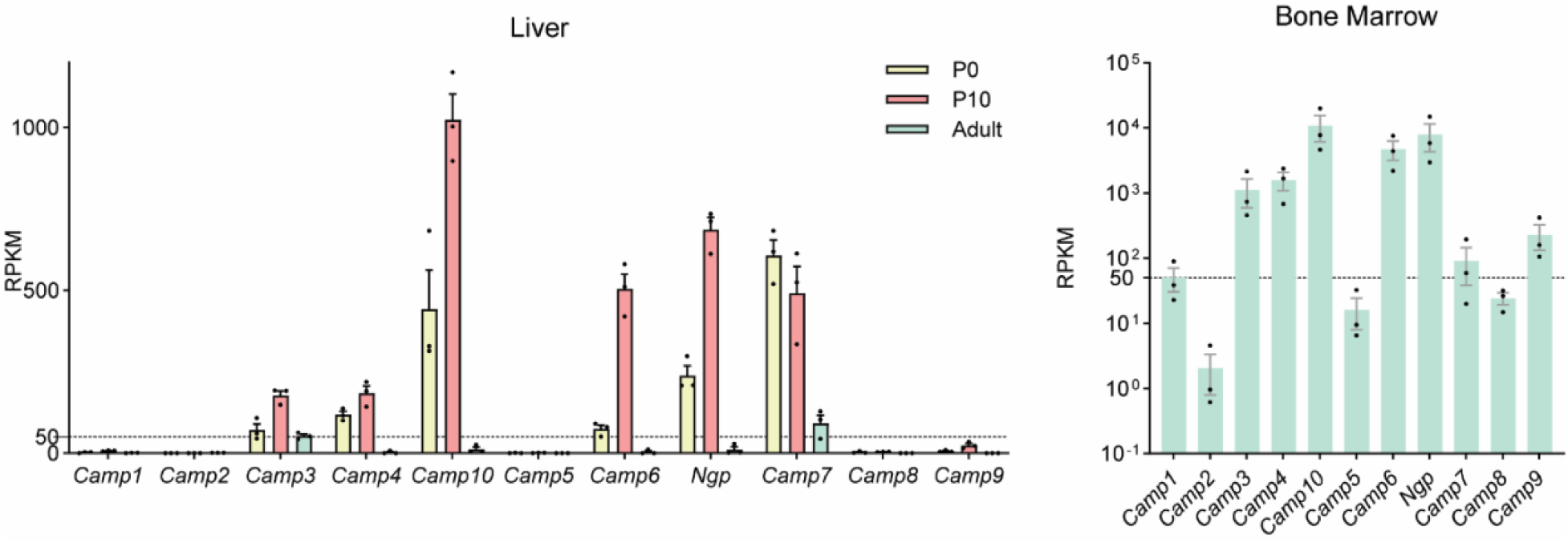
Bar graphs showing the expression of cathelicidin genes. Y-axis represents the Reads Per Kilobase per Million mapped reads (RPKM) values for liver and bone marrow tissue. Liver samples were profiled at various time points. Error bars represent standard error of the mean. Dotted lines denote RPKM of 50.

**Fig. S7.**
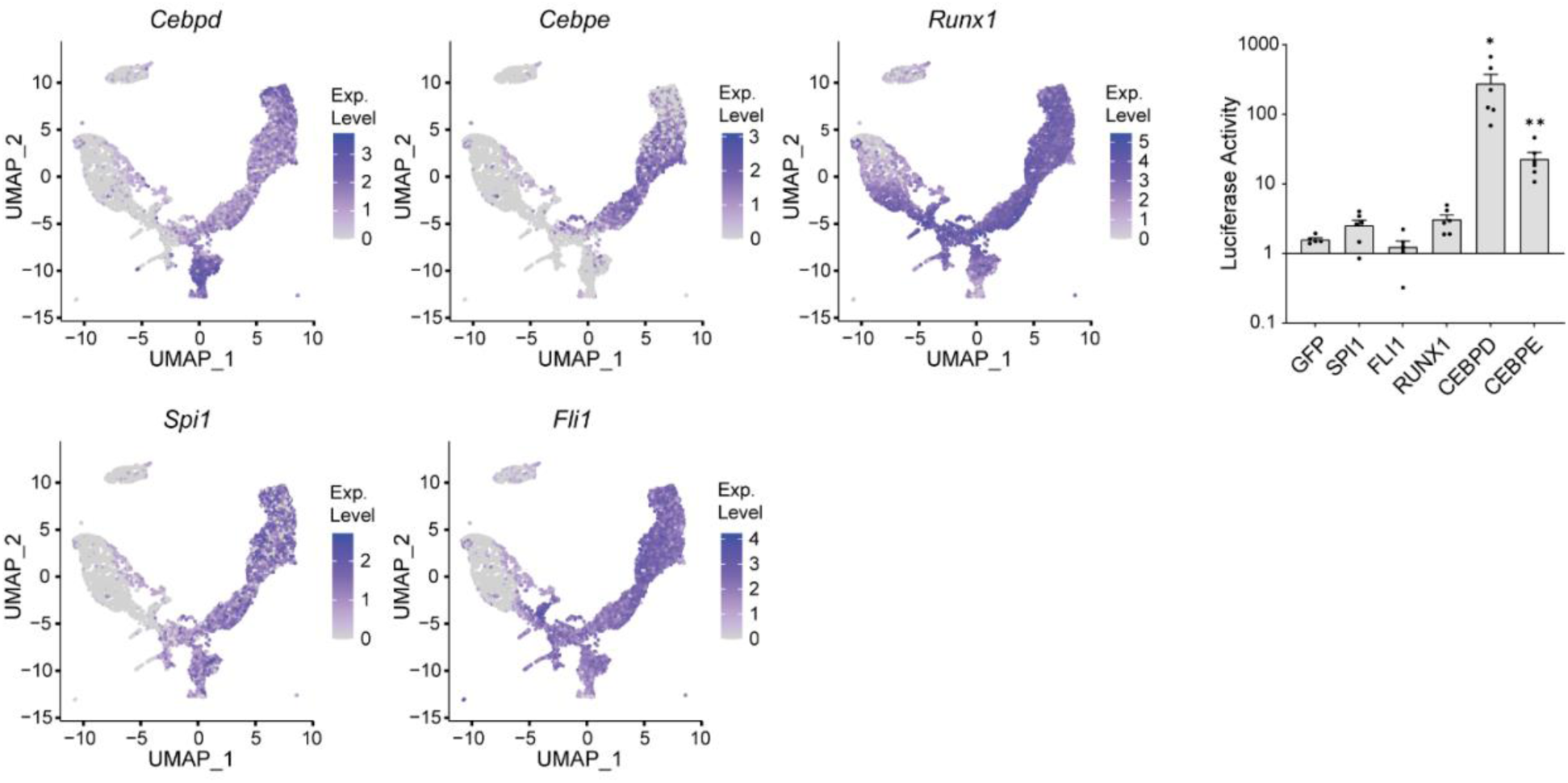
UMAP plots displaying the expression candidate transcription factors (left) and bar graphs showing the luciferase activity of the *Camp3* enhancer (right). Y-axis of the bar graphs represents the ratio of luminescence measured from the experimental reporter to the control reporter. Statistical significance was assessed using an unpaired t-test (n=5-6; *: p<0.05, **: p<0.01; Error bars correspond to standard error of the mean. Significance was recorded only when both biological replicates show significant upregulation compared to the GFP control).

**Fig. S8.**
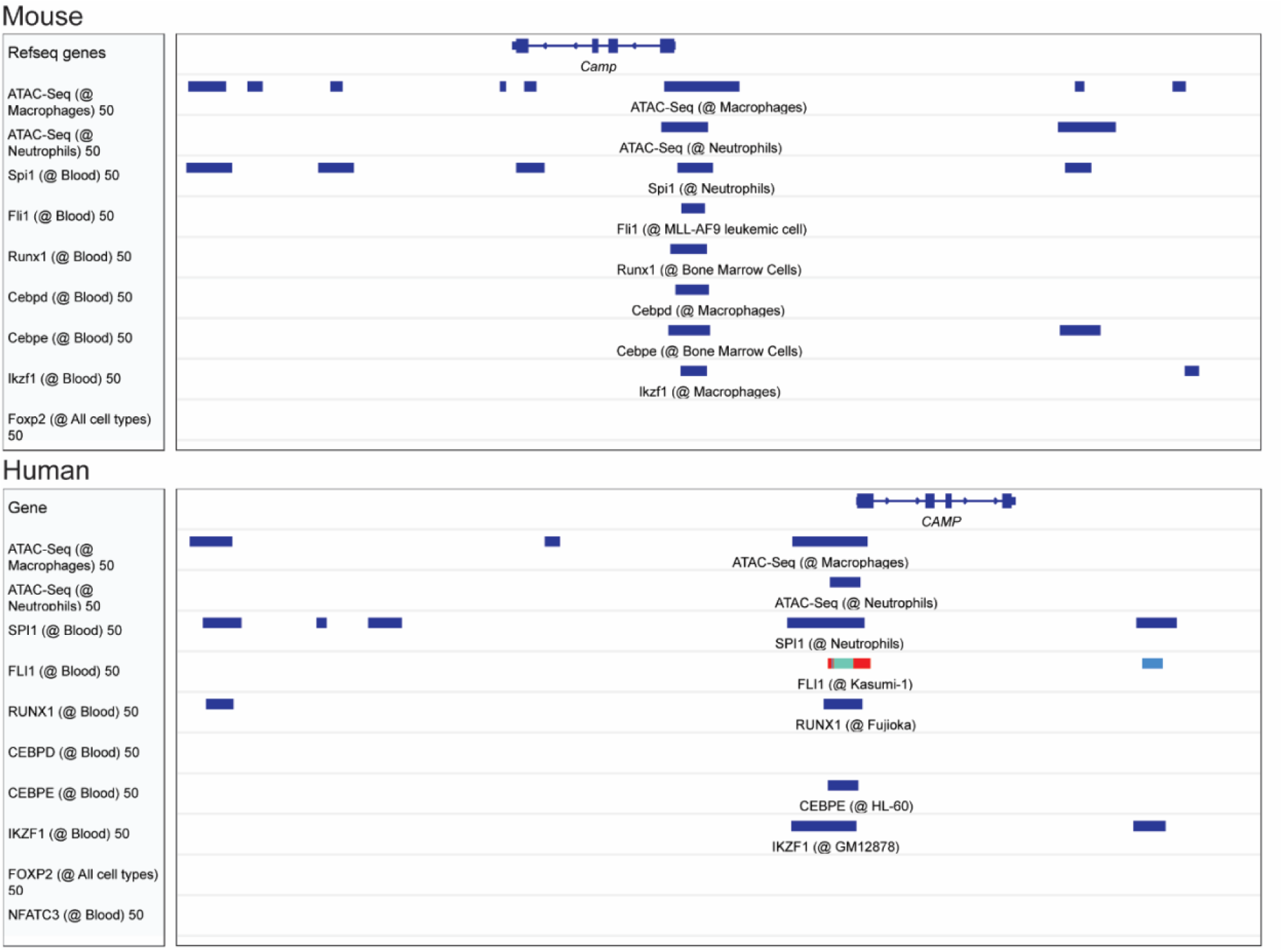
IGV snapshots showing ChIP-seq data of candidate transcription factors. Shown are data for mouse (top) and human (bottom). FOXP2 and NFATC3 were not known to bind to cathelicidin locus.

**Fig. S9.**
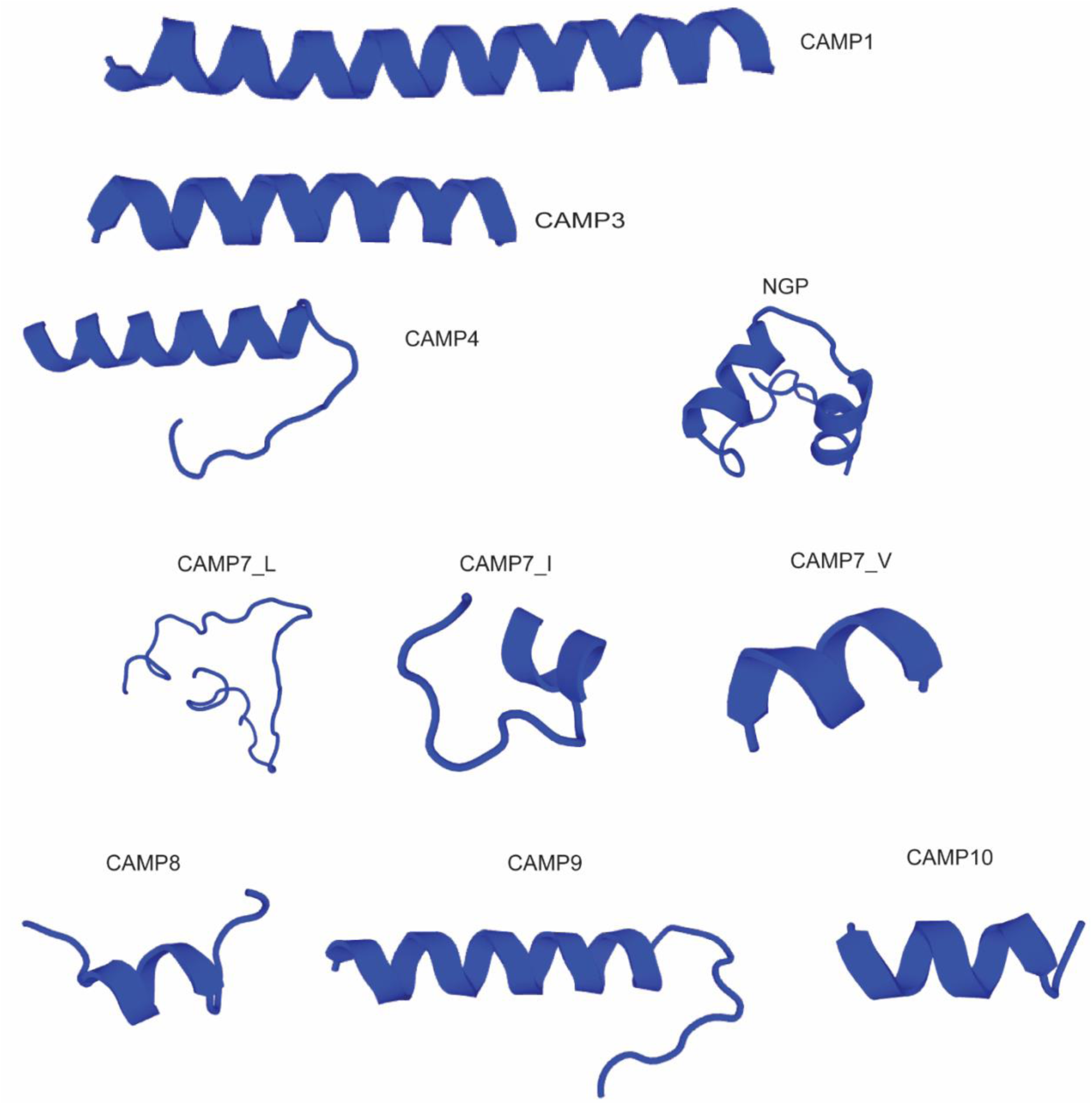
Predicted mature peptide structures of sugar glider cathelicidins. Secondary structures of synthesized peptides were predicted with PEP-FOLD3. Most peptides show canonical alpha-helix.

**Fig. S10.**
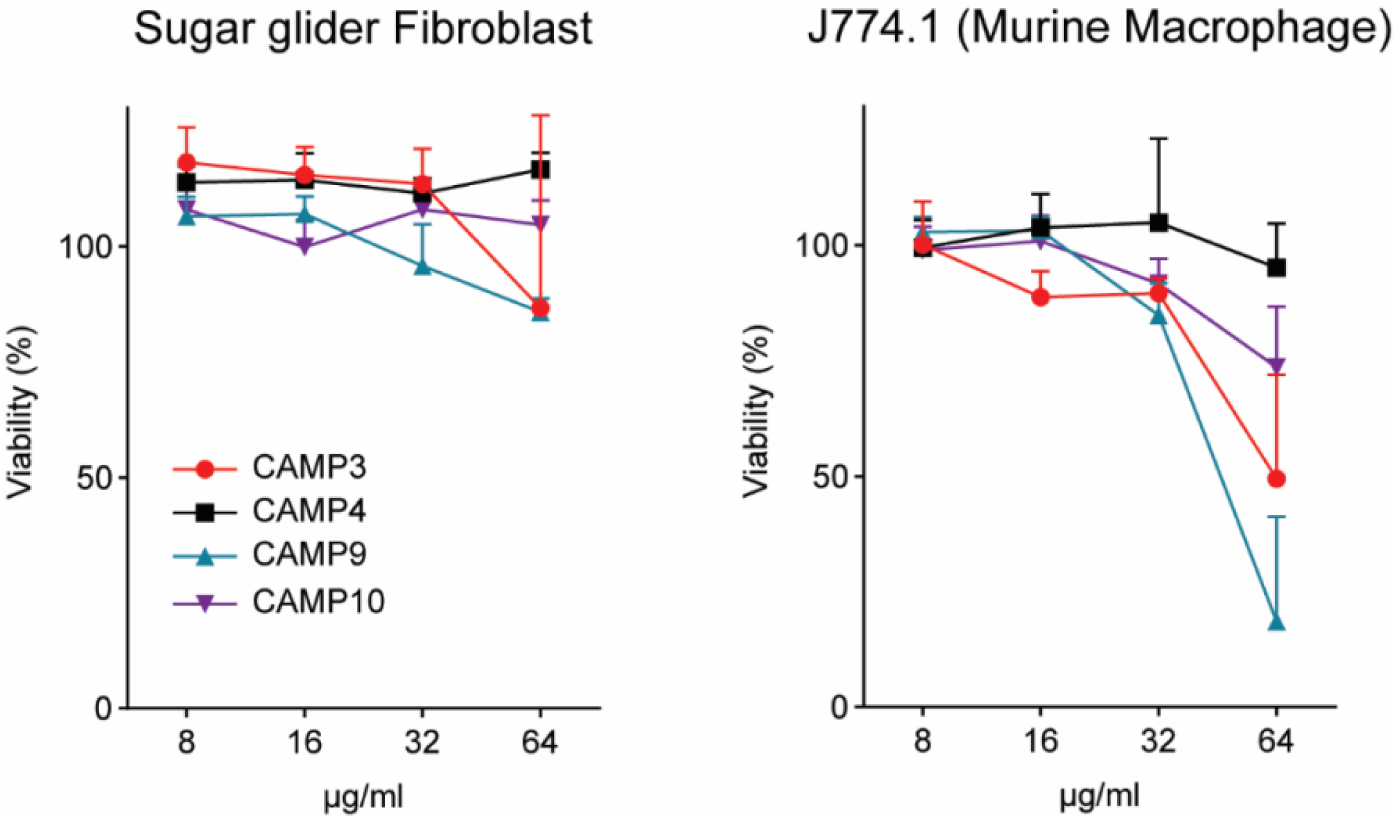
Cell viability assays. Shown is the percentage of viable sugar glider fibroblasts (left panel) and murine macrophages (right panel) after incubation with different concentrations of sugar glider cathelicidins.

**Fig. S11.**
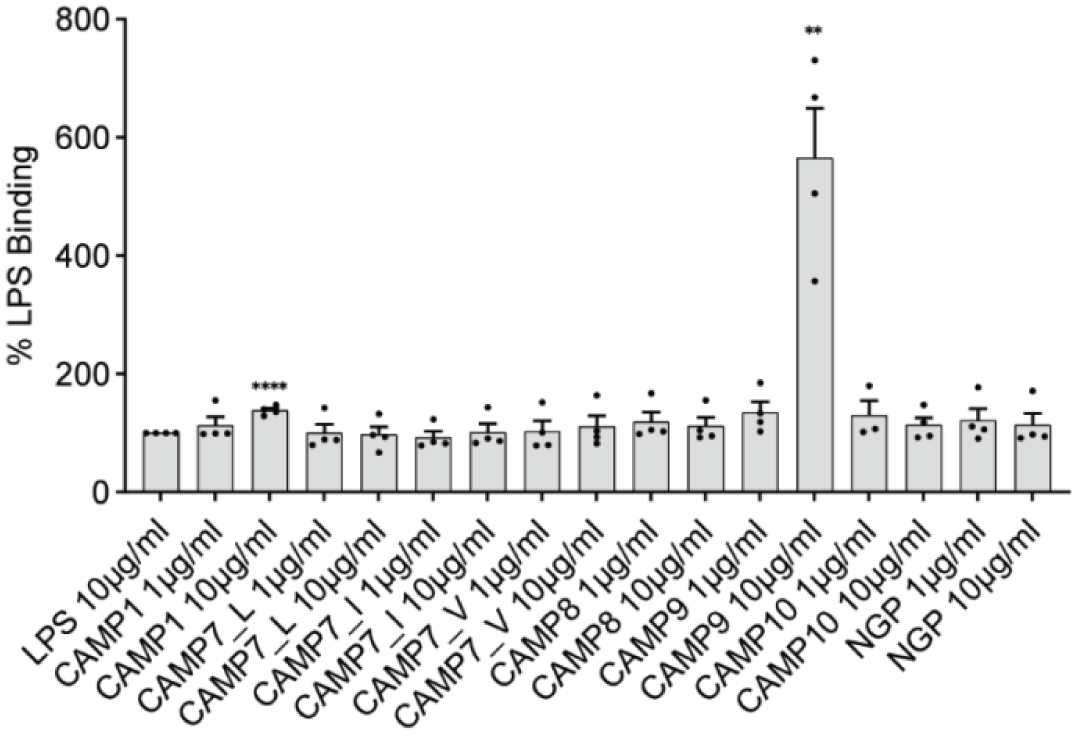
Bar graphs showing the result of LPS binding assay. Fluorescence intensities of J774A.1 cells treated with LPS-FITC were measured upon co-incubation with peptides or PBS control. The percentage of the median fluorescence intensity relative to the control is shown in the y-axis. Statistical significance was assessed using an unpaired *t*-test (n=4; **: p<0.01, ****:p<0.0001).

**Fig. S12.**
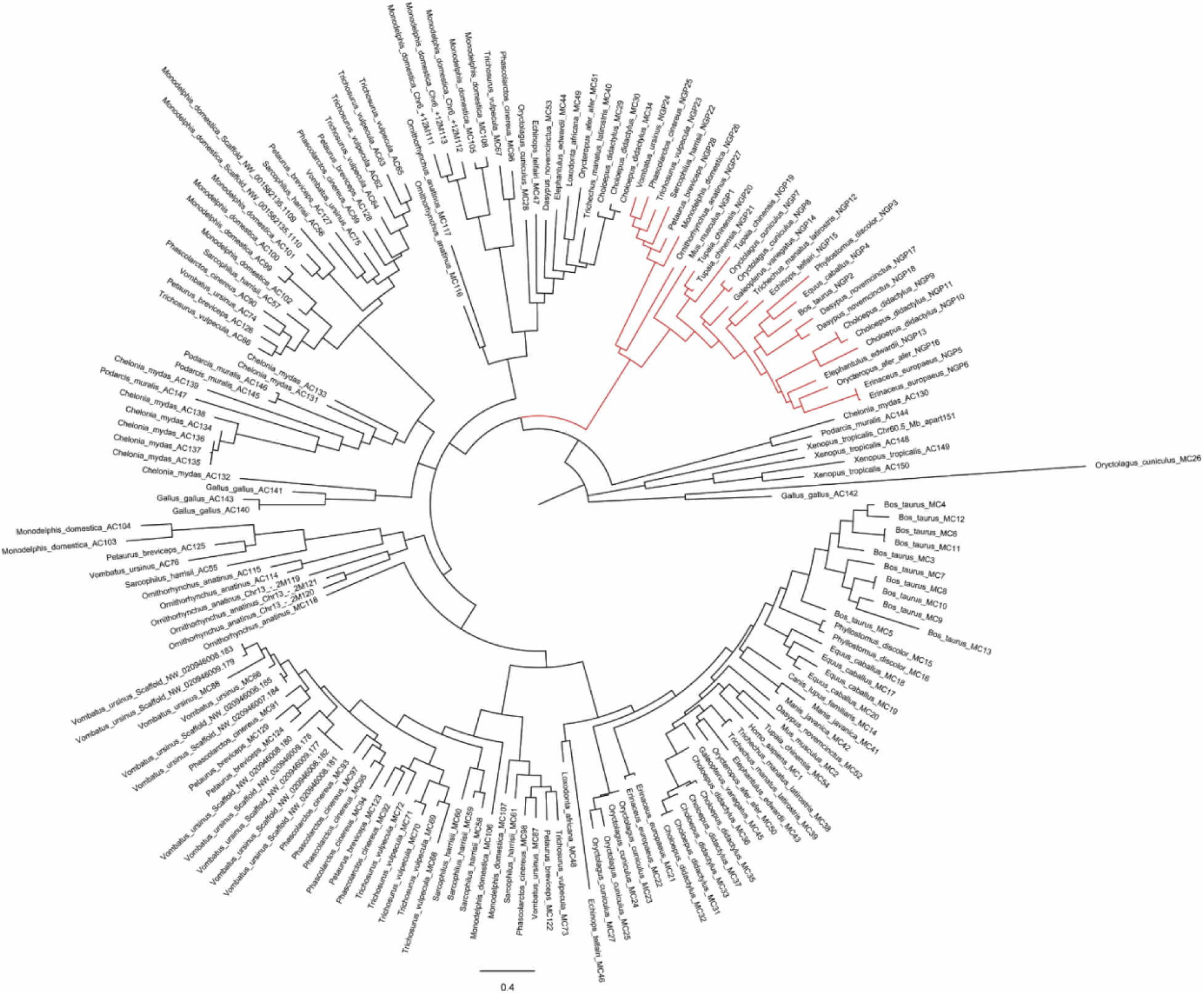
Maximum likelihood tree of 151 tetrapod cathelicidin genes and 28 mammalian NGP genes. All NGP genes form a monophyletic clade, indicating that NGP is a separate gene derived from cathelicidins (red). AC: ancestral cluster; MC: mammalian cluster.

**Table S1.**
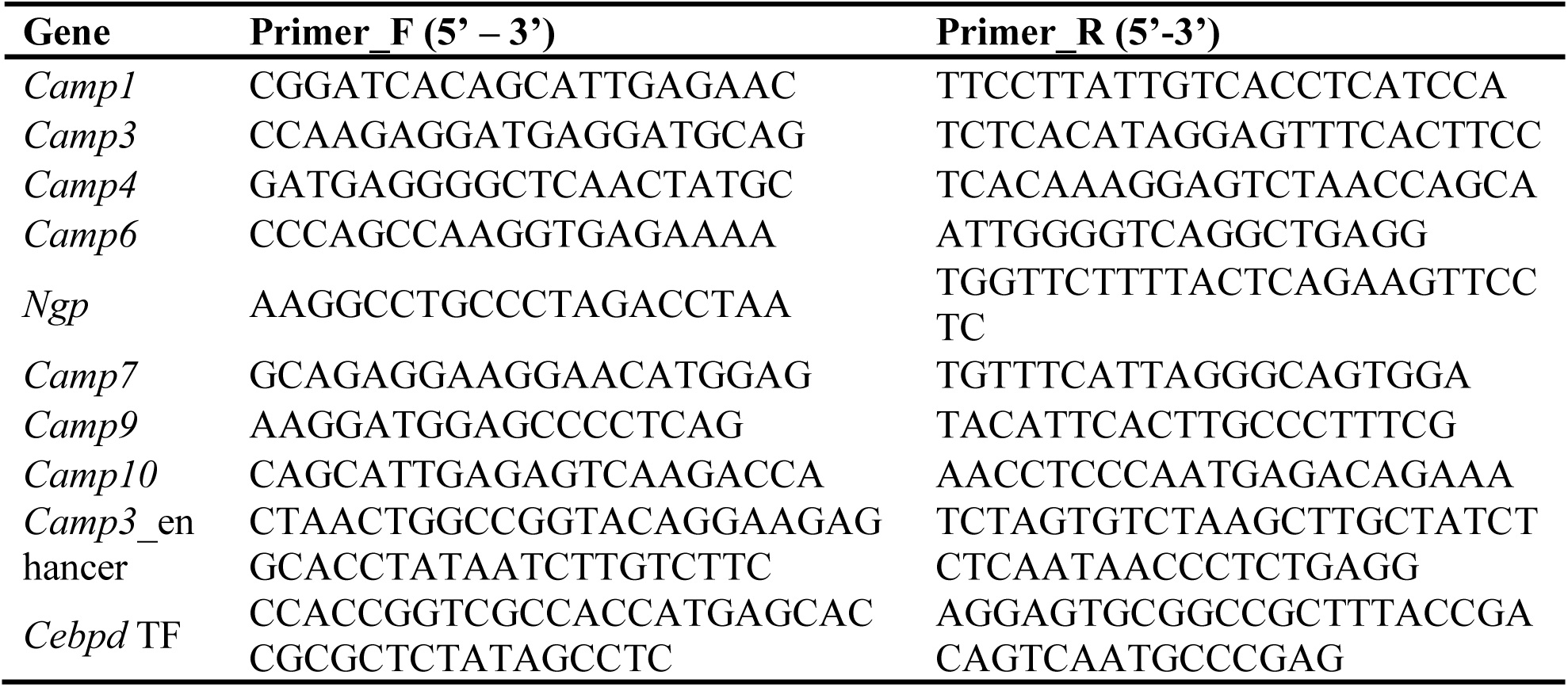
Primers used for annotation and cloning.

**Table S2.**
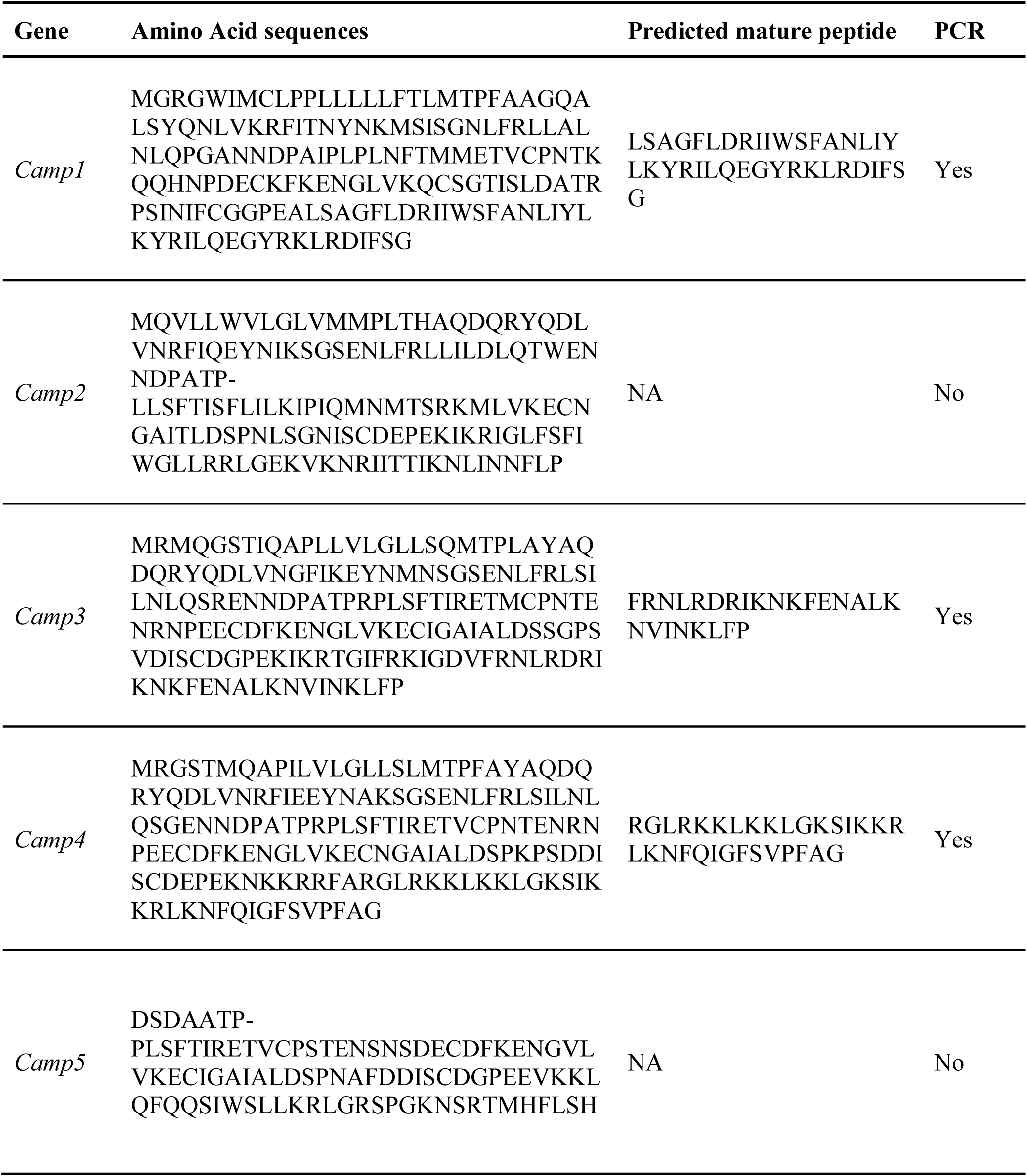
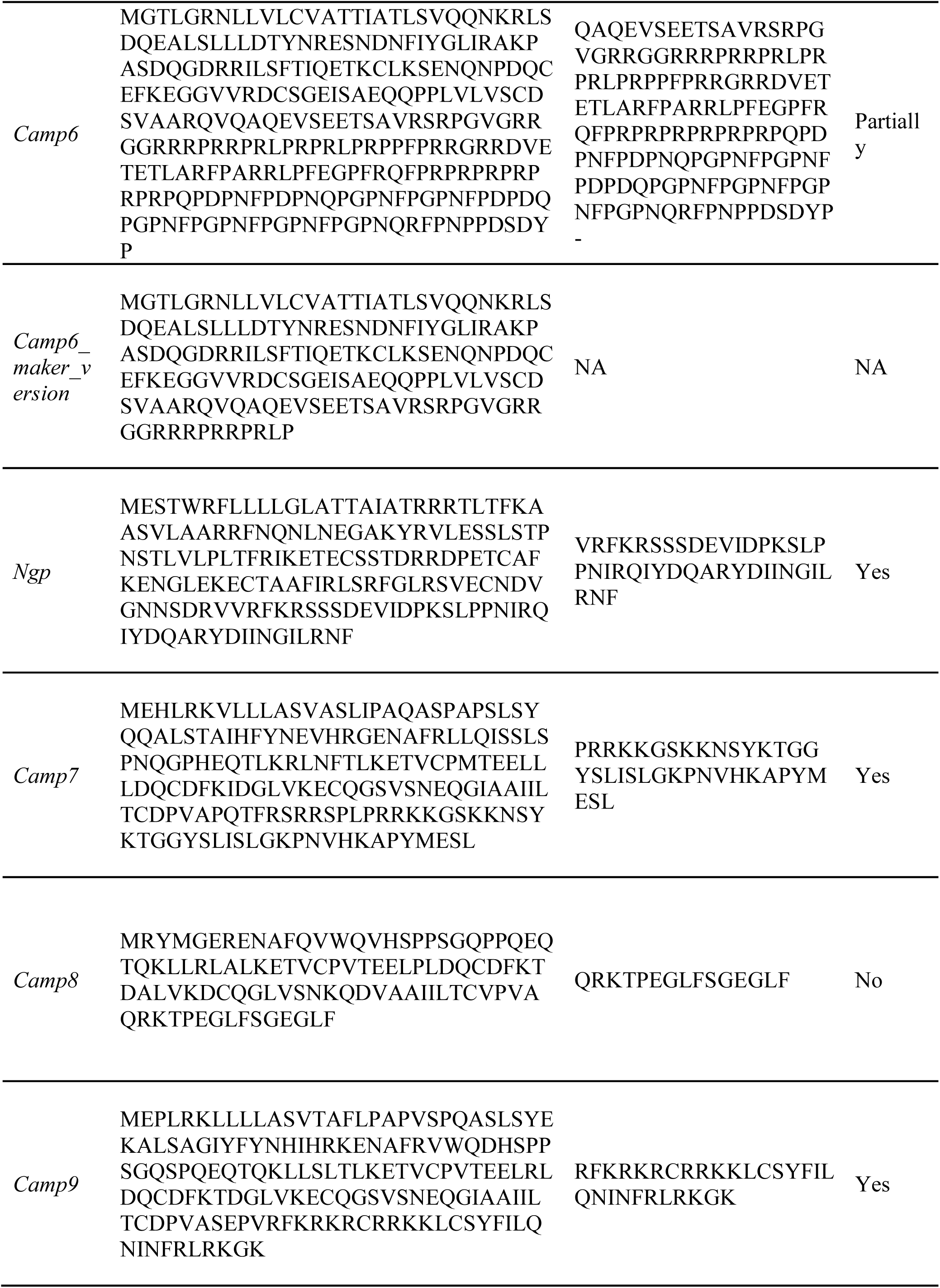
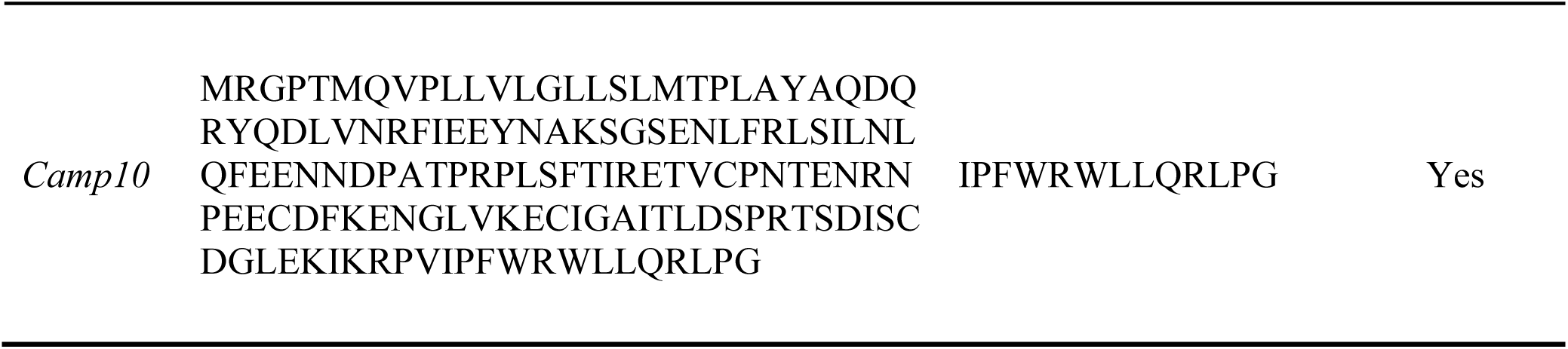
Cathelicidin sequence information. Amino acid sequences, predicted mature peptide sequences, and whether full length sequences are validated by Sanger sequencing are shown.

**Table S3.** Pairwise sequence identity matrix of cathelicidin amino acid sequences.

| Gene | CAMP3 | CAMP4 | CAMP10 | CAMP6 | CAMP7 | CAMP8 | CAMP9 |
| --- | --- | --- | --- | --- | --- | --- | --- |
| CAMP1 | 46.15 | 48.26 | 47.56 | 18.02 | 22.93 | 20.79 | 29.65 |
| CAMP3 |  | 69.19 | 69.7 | 19.27 | 25.52 | 20 | 26.78 |
| CAMP4 |  |  | 70.91 | 19.85 | 28.8 | 18.45 | 29.71 |
| CAMP10 |  |  |  | 17.65 | 27.22 | 20 | 28.41 |
| CAMP6 |  |  |  |  | 20.15 | 11.11 | 19.78 |
| CAMP7 |  |  |  |  |  | 34.3 | 56.14 |
| CAMP8 |  |  |  |  |  |  | 43.31 |

**Table S4.** Statistical significance of the luciferase assays.

|  | TF | Rep.1 | Rep.2 |
| --- | --- | --- | --- |
| Camp3_enhancer | *Spi1* | NS | 0.0332 |
|  | *Fli1* | NS | 0.0025 |
|  | *Runx1* | 0.0248 | NS |
|  | *Cebpd* | 0.0304 | <0.0001 |
|  | *Cebpe* | 0.0045 | 0.0003 |

**Table S5.**
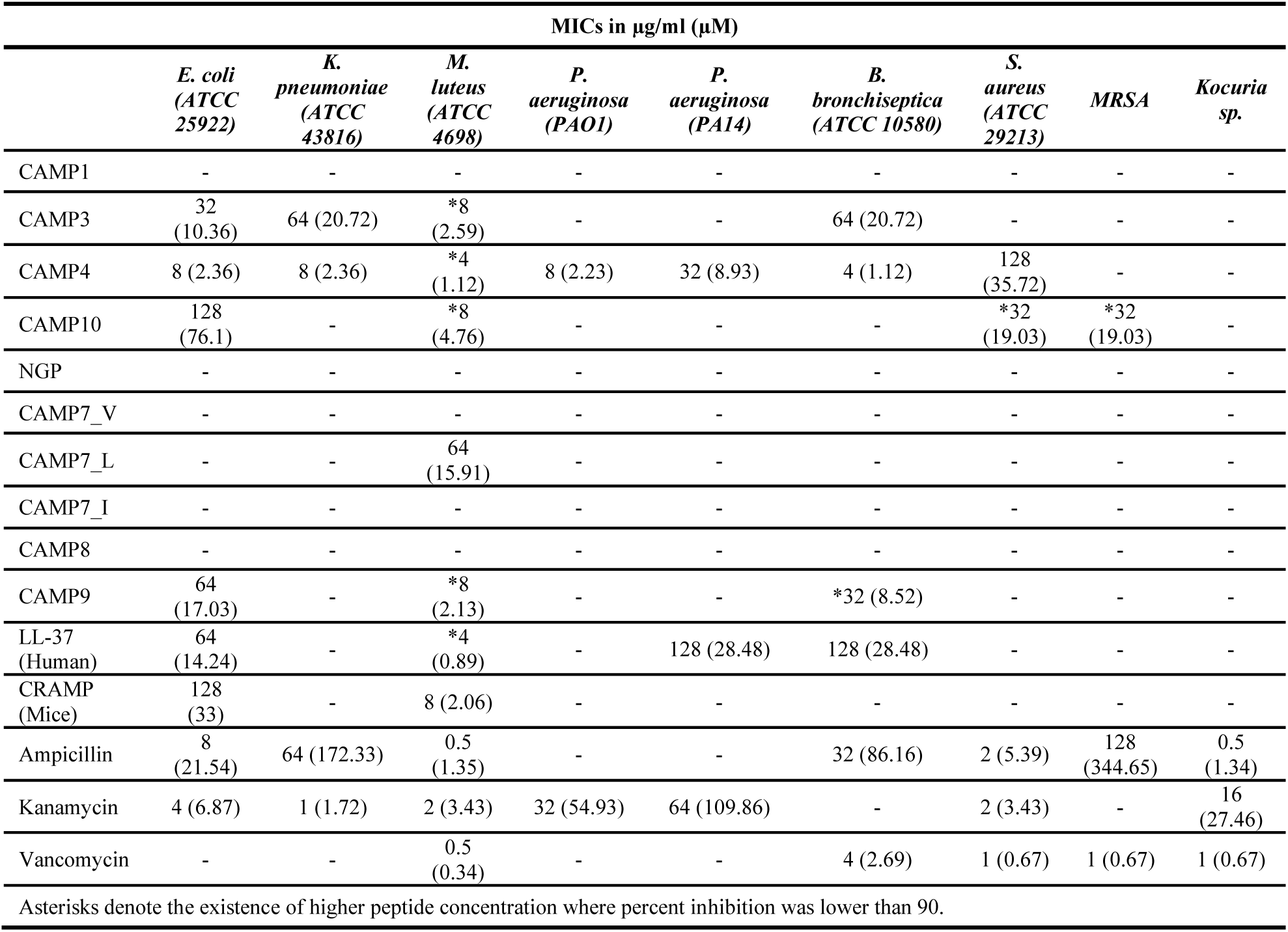
Minimal inhibitory concentrations (MICs) of synthesized peptides and control antibiotics.

**Table S6.**
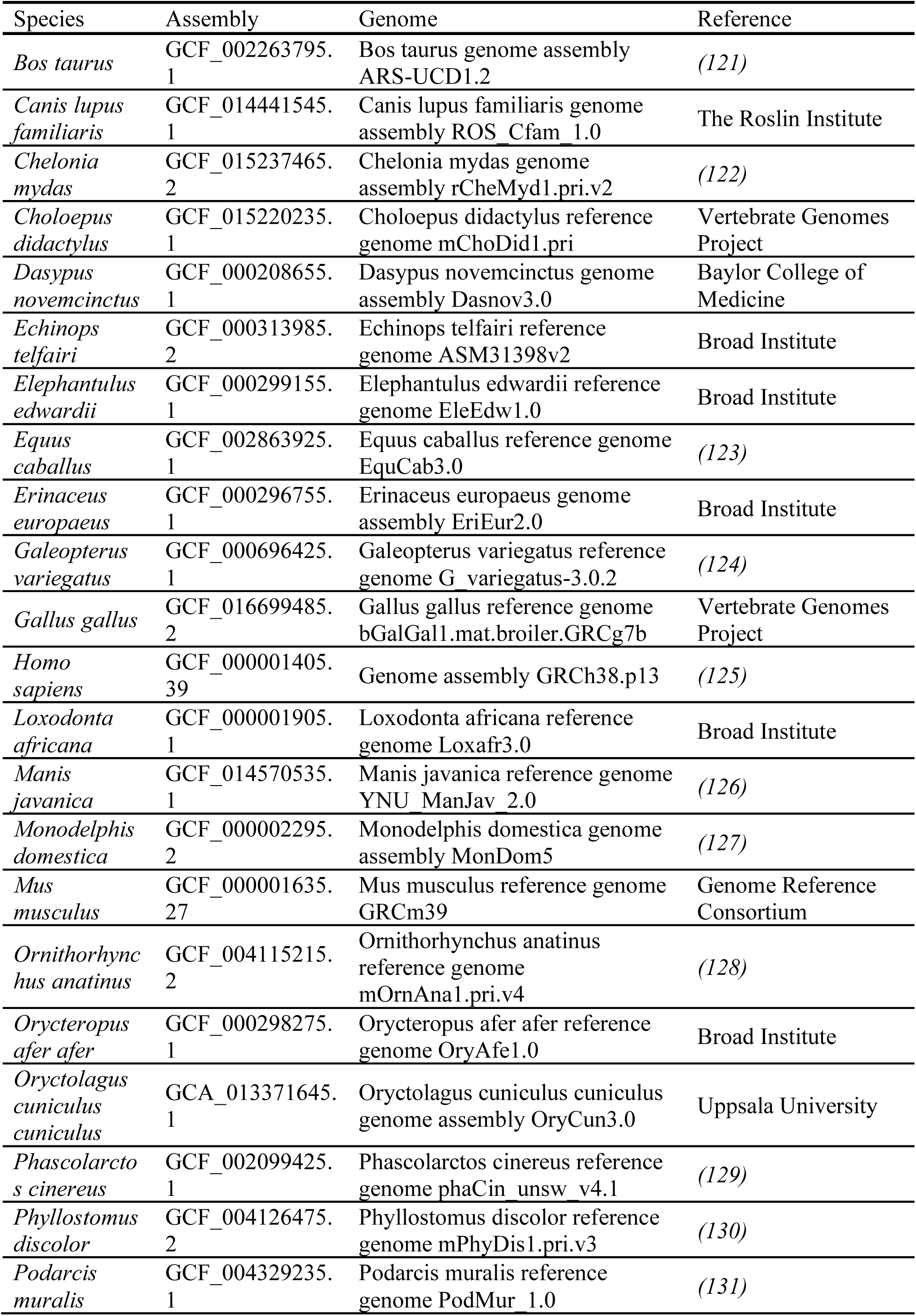
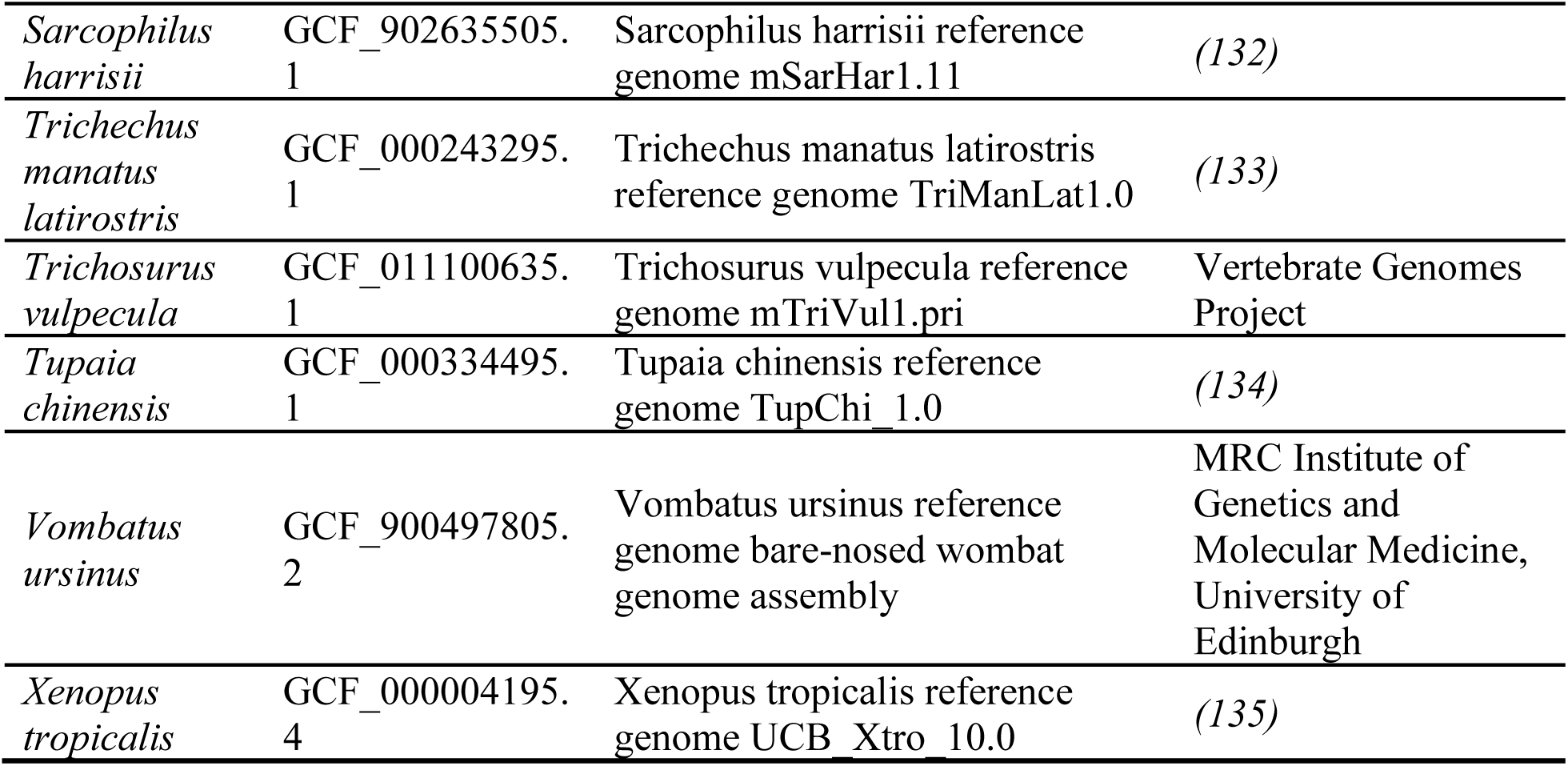
Genome information of 28 additional species used for syntenic analysis.

**Table S7.** Blast results of amino acid sequences of 4 representative mammalian *Ngp* against non-mammalian tetrapods.

| Query Species | Query gene | Subject species | Subject top hit |
| --- | --- | --- | --- |
| *Mus Musculus* | *Ngp* | Gallus gallus (taxid:9031) | cathelicidin-2 precursor |
| *Mus Musculus* | *Ngp* | Chelonia mydas (taxid:8469) | cathelicidin-related peptide Oh-Cath |
| *Mus Musculus* | *Ngp* | Podarcis muralis (taxid:64176) | cathelicidin-related peptide Oh-Cath-like |
| *Mus Musculus* | *Ngp* | Xenopus tropicalis (taxid:8364) | protegrin-3-like |
| *Trichosurus vulpecula* | *Ngp* | Gallus gallus (taxid:9031) | cathelicidin-2 precursor |
| *Trichosurus vulpecula* | *Ngp* | Chelonia mydas (taxid:8469) | uncharacterized protein LOC119565442 |
| *Trichosurus vulpecula* | *Ngp* | Podarcis muralis (taxid:64176) | uncharacterized protein LOC114607696 |
| *Trichosurus vulpecula* | *Ngp* | Xenopus tropicalis (taxid:8364) | protegrin-3-like |
| *Ornithorhynchus anatinus* | *Ngp* | Gallus gallus (taxid:9031) | cathelicidin-2 precursor |
| *Ornithorhynchus anatinus* | *Ngp* | Chelonia mydas (taxid:8469) | cathelicidin-related peptide Oh-Cath |
| *Ornithorhynchus anatinus* | *Ngp* | Podarcis muralis (taxid:64176) | cathelicidin-related peptide Oh-Cath-like |
| *Ornithorhynchus anatinus* | *Ngp* | Xenopus tropicalis (taxid:8364) | cathelicidin antimicrobial peptide |

**Data table S1. (separate file)**

596 genes differentially expressed genes in the comparison between neutrophil lineage cells and the rest of the liver cells (p-value < 0.0001, Log2Fold > 1).

**Data table S2. (separate file)**

Amino acid sequences of 79 mammalian cathelicidin and 26 *Ngp* sequences used for Maker3 annotation.

**Data table S3. (separate file)**

Results of FIMO (Find Individual Motif Occurences) motif scans.

**Data table S4. (separate file)**

Amino acid sequences of 151 tetrapod cathelicidin and 28 mammalian *Ngp* genes used for the construction of a phylogenetic tree.

## References

1. M. E. Perez-Muñoz, M.-C. Arrieta, A. E. Ramer-Tait, J. Walter, A critical assessment of the “sterile womb” and “in utero colonization” hypotheses: implications for research on the pioneer infant microbiome. Microbiome 5, 48 (2017).

2. M. C. de Goffau, S. Lager, U. Sovio, F. Gaccioli, E. Cook, S. J. Peacock, J. Parkhill, D. S. Charnock-Jones, G. C. S. Smith, Human placenta has no microbiome but can contain potential pathogens. Nature 572, 329–334 (2019).

3. A. A. Kuperman, A. Zimmerman, S. Hamadia, O. Ziv, V. Gurevich, B. Fichtman, N. Gavert, R. Straussman, H. Rechnitzer, M. Barzilay, S. Shvalb, J. Bornstein, I. Ben-Shachar, S. Yagel, I. Haviv, O. Koren, Deep microbial analysis of multiple placentas shows no evidence for a placental microbiome. BJOG 127, 159–169 (2020).

4. J. E. Lawn, S. Cousens, J. Zupan, Lancet Neonatal Survival Steering Team, 4 million neonatal deaths: when? Where? Why? Lancet 365, 891–900 (2005).

5. K. Sands, O. B. Spiller, K. Thomson, E. A. R. Portal, K. C. Iregbu, T. R. Walsh, Early-Onset Neonatal Sepsis in Low- and Middle-Income Countries: Current Challenges and Future Opportunities. Infect. Drug Resist. 15, 933–946 (2022).

6. A. Cassini, B. Allegranzi, C. Fleischmann-Struzek, T. Kortz, R. Markwart, H. Saito, M. Bonet, V. Brizuela, H. Mehrtash, Ö. Tuncalp Mingard, A.-B. Moller, O. Lincetto, S. Bellizzi, Y. Bin Nasr, B. Cao, R. Jakob, N. Tomaska, S. Eremin, P. Lambach, F. Moussy, “Global Report on the epidemiology and burden on sepsis: current evidence, identifying gaps and future directions” in Global Report on the Epidemiology and Burden on Sepsis: Current Evidence, Identifying Gaps and Future Directions. (World Health Organization, Geneva, 2020), p. 56.

7. C. C. L. Chase, D. J. Hurley, A. J. Reber, Neonatal immune development in the calf and its impact on vaccine response. Vet. Clin. North Am. Food Anim. Pract. 24, 87–104 (2008).

8. A. C. Jones, G. C. Burdge, J. A. Warner, A. D. Postle, J. O. Warner, Ontogeny of circulating leucocytes in the fetal guinea pig. Biol. Neonate 70, 108–115 (1996).

9. M. P. Holsapple, L. J. West, K. S. Landreth, Species comparison of anatomical and functional immune system development. Birth Defects Res. B Dev. Reprod. Toxicol. 68, 321– 334 (2003).

10. S. D. Holladay, R. J. Smialowicz, Development of the murine and human immune system: differential effects of immunotoxicants depend on time of exposure. Environ. Health Perspect. 108 **Suppl 3**, 463–473 (2000).

11. Z.-X. Luo, C.-X. Yuan, Q.-J. Meng, Q. Ji, A Jurassic eutherian mammal and divergence of marsupials and placentals. Nature 476, 442–445 (2011).

12. M. dos Reis, J. Inoue, M. Hasegawa, R. J. Asher, P. C. J. Donoghue, Z. Yang, Phylogenomic datasets provide both precision and accuracy in estimating the timescale of placental mammal phylogeny. Proc. Biol. Sci. 279, 3491–3500 (2012).

13. C. R. Borthwick, L. J. Young, J. M. Old, The development of the immune tissues in marsupial pouch young. J. Morphol. 275, 822–839 (2014).

14. J. M. Old, Haematopoiesis in Marsupials. Dev. Comp. Immunol. 58, 40–46 (2016).

15. J. E. Deakin, D. W. Cooper, Characterisation of and immunity to the aerobic bacteria found in the pouch of the brushtail possum Trichosurus vulpecula. Comp. Immunol. Microbiol. Infect. Dis. 27, 33–46 (2004).

16. R. Osawa, W. H. Blanshard, P. G. O’Callaghan, Microflora of the pouch of the koala (Phascolarctos cinereus). J. Wildl. Dis. 28, 276–280 (1992).

17. Y. Cheng, K. Belov, Antimicrobial Protection of Marsupial Pouch Young. Front. Microbiol. 8, 354 (2017).

18. K. M. Morris, D. O’Meally, T. Zaw, X. Song, A. Gillett, M. P. Molloy, A. Polkinghorne, K. Belov, Characterisation of the immune compounds in koala milk using a combined transcriptomic and proteomic approach. Sci. Rep. 6, 35011 (2016).

19. K. Ambatipudi, J. Joss, M. Raftery, E. Deane, A proteomic approach to analysis of antimicrobial activity in marsupial pouch secretions. Dev. Comp. Immunol. 32, 108–120 (2008).

20. K. Ambatipudi, J. Joss, E. Deane, A comparative proteomic analysis of skin secretions of the tammar wallaby (Macropus eugenii) and the wombat (Vombatus ursinus). Comp. Biochem. Physiol. Part D Genomics Proteomics 2, 322–331 (2007).

21. G. Bobek, E. M. Deane, Possible antimicrobial compounds from the pouch of the koala, Phascolarctos cinereus. Lett. Pept. Sci. 8, 133–137 (2001).

22. E. Peel, Y. Cheng, J. T. Djordjevic, S. Fox, T. C. Sorrell, K. Belov, Cathelicidins in the Tasmanian devil (Sarcophilus harrisii). Sci. Rep. 6, 35019 (2016).

23. C. Petrohilos, A. Patchett, C. J. Hogg, K. Belov, E. Peel, Tasmanian devil cathelicidins exhibit anticancer activity against Devil Facial Tumour Disease (DFTD) cells. Sci. Rep. 13, 12698 (2023).

24. R. V. Hewavisenti, K. M. Morris, D. O’Meally, Y. Cheng, A. T. Papenfuss, K. Belov, The identification of immune genes in the milk transcriptome of the Tasmanian devil (Sarcophilus harrisii). PeerJ 4, e1569 (2016).

25. C. R. Borthwick, L. J. Young, J. M. Old, An Examination of the Development and Localization of Key Immune Cells in Developing Pouch Young of the Red-Tailed Phascogale (Phascogale calura). Anat. Rec. 302, 1985–2002 (2019).

26. J. M. Old, E. M. Deane, Immunohistochemistry of the lymphoid tissues of the tammar wallaby, Macropus eugenii. J. Anat. 201, 257–266 (2002).

27. S. W. Hemsley, P. J. Canfield, A. J. Husband, Immunohistological staining of lymphoid tissue in four Australian marsupial species using species cross-reactive antibodies. Immunol. Cell Biol. 73, 321–325 (1995).

28. S. W. Hemsley, P. J. Canfield, A. J. Husband, Histological and immunohistological investigation of alimentary tract lymphoid tissue in the koala (Phascolarctos cinereus), brushtail possum (Trichosurus vulpecula) and ringtail possum (Pseudocheirus peregrinus). J. Anat. 188 **(Pt** **2**), 279–288 (1996).

29. L. J. Howson, K. M. Morris, T. Kobayashi, C. Tovar, A. Kreiss, A. T. Papenfuss, L. Corcoran, K. Belov, G. M. Woods, Identification of dendritic cells, B cell and T cell subsets in Tasmanian devil lymphoid tissue; evidence for poor immune cell infiltration into devil facial tumors. Anat. Rec. 297, 925–938 (2014).

30. L. G. Duncan, S. V. Nair, E. M. Deane, Immunohistochemical localization of T-lymphocyte subsets in the developing lymphoid tissues of the tammar wallaby (Macropus eugenii). Dev. Comp. Immunol. 38, 475–486 (2012).

31. G. L. Burn, A. Foti, G. Marsman, D. F. Patel, A. Zychlinsky, The Neutrophil. Immunity 54, 1377–1391 (2021).

32. P. A. Cisternas, P. J. Armati, Development of the thymus, spleen, lymph nodes and liver in the marsupial, Isoodon macrourus (Northern brown bandicoot, Peramelidae). Anat. Embryol. 200, 433–443 (1999).

33. J. M. Old, L. Selwood, E. M. Deane, A developmental investigation of the liver, bone marrow and spleen of the stripe-faced dunnart (Sminthopsis macroura). Dev. Comp. Immunol. 28, 347–355 (2004).

34. A. E. Shinnar, K. L. Butler, H. J. Park, Cathelicidin family of antimicrobial peptides: proteolytic processing and protease resistance. Bioorg. Chem. 31, 425–436 (2003).

35. E. M. Kościuczuk, P. Lisowski, J. Jarczak, N. Strzałkowska, A. Jóźwik, J. Horbańczuk, J. Krzyżewski, L. Zwierzchowski, E. Bagnicka, Cathelicidins: family of antimicrobial peptides. A review. Mol. Biol. Rep. 39, 10957–10970 (2012).

36. M. A. Alford, B. Baquir, F. L. Santana, E. F. Haney, R. E. W. Hancock, Cathelicidin Host Defense Peptides and Inflammatory Signaling: Striking a Balance. Front. Microbiol. 11, 1902 (2020).

37. K. A. Daly, M. R. Digby, C. Lefévre, K. R. Nicholas, E. M. Deane, P. Williamson, Identification, characterization and expression of cathelicidin in the pouch young of tammar wallaby (Macropus eugenii). Comp. Biochem. Physiol. B Biochem. Mol. Biol. 149, 524–533 (2008).

38. E. Peel, Y. Cheng, J. T. Djordjevic, D. O’Meally, M. Thomas, M. Kuhn, T. C. Sorrell, W. M. Huston, K. Belov, Koala cathelicidin PhciCath5 has antimicrobial activity, including against Chlamydia pecorum. PLoS One 16, e0249658 (2021).

39. E. Peel, Y. Cheng, J. T. Djordjevic, M. Kuhn, T. Sorrell, K. Belov, Marsupial and monotreme cathelicidins display antimicrobial activity, including against methicillin-resistant Staphylococcus aureus. Microbiology 163, 1457–1465 (2017).

40. K. Belov, C. E. Sanderson, J. E. Deakin, E. S. W. Wong, D. Assange, K. A. McColl, A. Gout, B. de Bono, A. D. Barrow, T. P. Speed, J. Trowsdale, A. T. Papenfuss, Characterization of the opossum immune genome provides insights into the evolution of the mammalian immune system. Genome Res. 17, 982–991 (2007).

41. J. Wang, E. S. W. Wong, J. C. Whitley, J. Li, J. M. Stringer, K. R. Short, M. B. Renfree, K. Belov, B. G. Cocks, Ancient antimicrobial peptides kill antibiotic-resistant pathogens: Australian mammals provide new options. PLoS One 6, e24030 (2011).

42. H.-S. Cho, J. Yum, A. Larivière, N. Lévêque, Q. V. C. Le, B. Ahn, H. Jeon, K. Hong, N. Soundrarajan, J.-H. Kim, C. Bodet, C. Park, Opossum Cathelicidins Exhibit Antimicrobial Activity Against a Broad Spectrum of Pathogens Including West Nile Virus. Front. Immunol. 11, 347 (2020).

43. R. L. Carman, J. M. Old, M. Baker, N. A. Jacques, E. M. Deane, Identification and expression of a novel marsupial cathelicidin from the tammar wallaby (Macropus eugenii). Vet. Immunol. Immunopathol. 127, 269–276 (2009).

44. C. Y. Feigin, J. A. Moreno, R. Ramos, S. A. Mereby, A. Alivisatos, W. Wang, R. van Amerongen, J. Camacho, J. J. Rasweiler 4th, R. R. Behringer, B. Ostrow, M. V. Plikus, R. Mallarino, Convergent deployment of ancestral functions during the evolution of mammalian flight membranes. Sci Adv 9, eade7511 (2023).

45. J. D. Buenrostro, P. G. Giresi, L. C. Zaba, H. Y. Chang, W. J. Greenleaf, Transposition of native chromatin for fast and sensitive epigenomic profiling of open chromatin, DNA-binding proteins and nucleosome position. Nat. Methods 10, 1213–1218 (2013).

46. M. R. Corces, A. E. Trevino, E. G. Hamilton, P. G. Greenside, N. A. Sinnott-Armstrong, S. Vesuna, A. T. Satpathy, A. J. Rubin, K. S. Montine, B. Wu, A. Kathiria, S. W. Cho, M. R. Mumbach, A. C. Carter, M. Kasowski, L. A. Orloff, V. I. Risca, A. Kundaje, P. A. Khavari, T. J. Montine, W. J. Greenleaf, H. Y. Chang, An improved ATAC-seq protocol reduces background and enables interrogation of frozen tissues. Nat. Methods 14, 959–962 (2017).

47. J. A. Moreno, O. Dudchenko, C. Y. Feigin, S. A. Mereby, Z. Chen, R. Ramos, A. A. Almet, H. Sen, B. J. Brack, M. R. Johnson, S. Li, W. Wang, J. M. Gaska, A. Ploss, D. Weisz, A. D. Omer, W. Yao, Z. Colaric, P. Kaur, J. S. Leger, Q. Nie, A. Mena, J. P. Flanagan, G. Keller, T. Sanger, B. Ostrow, M. V. Plikus, E. Z. Kvon, E. L. Aiden, R. Mallarino, Emx2 underlies the development and evolution of marsupial gliding membranes. Nature 629, 127–135 (2024).

48. V. Y. Goel, M. K. Huseyin, A. S. Hansen, Region Capture Micro-C reveals coalescence of enhancers and promoters into nested microcompartments. Nat. Genet. 55, 1048–1056 (2023).

49. O. E. Sørensen, P. Follin, A. H. Johnsen, J. Calafat, G. S. Tjabringa, P. S. Hiemstra, N. Borregaard, Human cathelicidin, hCAP-18, is processed to the antimicrobial peptide LL-37 by extracellular cleavage with proteinase 3. Blood 97, 3951–3959 (2001).

50. A. M. Cole, J. Shi, A. Ceccarelli, Y. H. Kim, A. Park, T. Ganz, Inhibition of neutrophil elastase prevents cathelicidin activation and impairs clearance of bacteria from wounds. Blood 97, 297–304 (2001).

51. M. Scocchi, B. Skerlavaj, D. Romeo, R. Gennaro, Proteolytic cleavage by neutrophil elastase converts inactive storage proforms to antibacterial bactenecins. Eur. J. Biochem. 209, 589– 595 (1992).

52. A. Panyutich, J. Shi, P. L. Boutz, C. Zhao, T. Ganz, Porcine polymorphonuclear leukocytes generate extracellular microbicidal activity by elastase-mediated activation of secreted proprotegrins. Infect. Immun. 65, 978–985 (1997).

53. R. L. Gallo, K. J. Kim, M. Bernfield, C. A. Kozak, M. Zanetti, L. Merluzzi, R. Gennaro, Identification of CRAMP, a cathelin-related antimicrobial peptide expressed in the embryonic and adult mouse. J. Biol. Chem. 272, 13088–13093 (1997).

54. B. Agerberth, H. Gunne, J. Odeberg, P. Kogner, H. G. Boman, G. H. Gudmundsson, FALL-39, a putative human peptide antibiotic, is cysteine-free and expressed in bone marrow and testis. Proc. Natl. Acad. Sci. U. S. A. 92, 195–199 (1995).

55. E. Sancho-Vaello, D. Gil-Carton, P. François, E.-J. Bonetti, M. Kreir, K. R. Pothula, U. Kleinekathöfer, K. Zeth, The structure of the antimicrobial human cathelicidin LL-37 shows oligomerization and channel formation in the presence of membrane mimics. Sci. Rep. 10, 17356 (2020).

56. M. Scocchi, I. Zelezetsky, M. Benincasa, R. Gennaro, A. Mazzoli, A. Tossi, Structural aspects and biological properties of the cathelicidin PMAP-36. FEBS J. 272, 4398–4406 (2005).

57. L. Boulos, M. Prévost, B. Barbeau, J. Coallier, R. Desjardins, LIVE/DEAD® BacLight^TM^: application of a new rapid staining method for direct enumeration of viable and total bacteria in drinking water. J. Microbiol. Methods 37, 77–86 (1999).

58. J. Koziel, D. Bryzek, A. Sroka, K. Maresz, I. Glowczyk, E. Bielecka, T. Kantyka, K. Pyrć, P. Svoboda, J. Pohl, J. Potempa, Citrullination alters immunomodulatory function of LL-37 essential for prevention of endotoxin-induced sepsis. J. Immunol. 192, 5363–5372 (2014).

59. F. Niyonsaba, K. Iwabuchi, A. Someya, M. Hirata, H. Matsuda, H. Ogawa, I. Nagaoka, A cathelicidin family of human antibacterial peptide LL-37 induces mast cell chemotaxis. Immunology 106, 20–26 (2002).

60. M. G. Scott, D. J. Davidson, M. R. Gold, D. Bowdish, R. E. W. Hancock, The human antimicrobial peptide LL-37 is a multifunctional modulator of innate immune responses. J. Immunol. 169, 3883–3891 (2002).

61. D. Yang, Q. Chen, A. P. Schmidt, G. M. Anderson, J. M. Wang, J. Wooters, J. J. Oppenheim, O. Chertov, Ll-37, the Neutrophil Granule–And Epithelial Cell–Derived Cathelicidin, Utilizes Formyl Peptide Receptor–Like 1 (Fprl1) as a Receptor to Chemoattract Human Peripheral Blood Neutrophils, Monocytes, and T Cells. J. Exp. Med. 192, 1069–1074 (2000).

62. A. M. van der Does, H. Beekhuizen, B. Ravensbergen, T. Vos, T. H. M. Ottenhoff, J. T. van Dissel, J. W. Drijfhout, P. S. Hiemstra, P. H. Nibbering, LL-37 directs macrophage differentiation toward macrophages with a proinflammatory signature. J. Immunol. 185, 1442–1449 (2010).

63. D. Minns, K. J. Smith, V. Alessandrini, G. Hardisty, L. Melrose, L. Jackson-Jones, A. S. MacDonald, D. J. Davidson, E. Gwyer Findlay, The neutrophil antimicrobial peptide cathelicidin promotes Th17 differentiation. Nat. Commun. 12, 1285 (2021).

64. N. J. Greenfield, Using circular dichroism spectra to estimate protein secondary structure. Nat. Protoc. 1, 2876–2890 (2006).

65. A. J. Lewis, C. W. Seymour, M. R. Rosengart, Current Murine Models of Sepsis. Surg. Infect. 17, 385–393 (2016).

66. A. Shrestha, D. Duwadi, J. Jukosky, S. N. Fiering, Cecropin-like antimicrobial peptide protects mice from lethal E.coli infection. PLoS One 14, e0220344 (2019).

67. K. Ferner, J. A. Schultz, U. Zeller, Comparative anatomy of neonates of the three major mammalian groups (monotremes, marsupials, placentals) and implications for the ancestral mammalian neonate morphotype. J. Anat. 231, 798–822 (2017).

68. H. Kiyonari, M. Kaneko, T. Abe, A. Shiraishi, R. Yoshimi, K.-I. Inoue, Y. Furuta, Targeted gene disruption in a marsupial, Monodelphis domestica, by CRISPR/Cas9 genome editing. Curr. Biol. 31, 3956–3963.e4 (2021).

69. Y. Hao, S. Hao, E. Andersen-Nissen, W. M. Mauck 3rd, S. Zheng, A. Butler, M. J. Lee, A. J. Wilk, C. Darby, M. Zager, P. Hoffman, M. Stoeckius, E. Papalexi, E. P. Mimitou, J. Jain, A. Srivastava, T. Stuart, L. M. Fleming, B. Yeung, A. J. Rogers, J. M. McElrath, C. A. Blish, R. Gottardo, P. Smibert, R. Satija, Integrated analysis of multimodal single-cell data. Cell 184, 3573–3587.e29 (2021).

70. L. V. Yang, R. H. Nicholson, J. Kaplan, A. Galy, L. Li, Hemogen is a novel nuclear factor specifically expressed in mouse hematopoietic development and its human homologue EDAG maps to chromosome 9q22, a region containing breakpoints of hematological neoplasms. Mech. Dev. 104, 105–111 (2001).

71. S. Elliott, A. M. Sinclair, The effect of erythropoietin on normal and neoplastic cells. Biologics 6, 163–189 (2012).

72. B. S. Garrison, A. P. Rybak, I. Beerman, B. Heesters, F. E. Mercier, D. T. Scadden, D. Bryder, R. Baron, D. J. Rossi, ZFP521 regulates murine hematopoietic stem cell function and facilitates MLL-AF9 leukemogenesis in mouse and human cells. Blood 130, 619–624 (2017).

73. D. B. AbuSamra, F. A. Aleisa, A. S. Al-Amoodi, H. M. Jalal Ahmed, C. J. Chin, A. F. Abuelela, P. Bergam, R. Sougrat, J. S. Merzaban, Not just a marker: CD34 on human hematopoietic stem/progenitor cells dominates vascular selectin binding along with CD44. Blood Adv 1, 2799–2816 (2017).

74. M. Evrard, I. W. H. Kwok, S. Z. Chong, K. W. W. Teng, E. Becht, J. Chen, J. L. Sieow, H. L. Penny, G. C. Ching, S. Devi, J. M. Adrover, J. L. Y. Li, K. H. Liong, L. Tan, Z. Poon, S. Foo, J. W. Chua, I.-H. Su, K. Balabanian, F. Bachelerie, S. K. Biswas, A. Larbi, W. Y. K. Hwang, V. Madan, H. P. Koeffler, S. C. Wong, E. W. Newell, A. Hidalgo, F. Ginhoux, L. G. Ng, Developmental Analysis of Bone Marrow Neutrophils Reveals Populations Specialized in Expansion, Trafficking, and Effector Functions. Immunity 48, 364–379.e8 (2018).

75. S. R. Horman, C. S. Velu, A. Chaubey, T. Bourdeau, J. Zhu, W. E. Paul, B. Gebelein, H. L. Grimes, Gfi1 integrates progenitor versus granulocytic transcriptional programming. Blood 113, 5466–5475 (2009).

76. X. Xie, Q. Shi, P. Wu, X. Zhang, H. Kambara, J. Su, H. Yu, S.-Y. Park, R. Guo, Q. Ren, S. Zhang, Y. Xu, L. E. Silberstein, T. Cheng, F. Ma, C. Li, H. R. Luo, Single-cell transcriptome profiling reveals neutrophil heterogeneity in homeostasis and infection. Nat. Immunol. 21, 1119–1133 (2020).

77. P. A. Szabo, H. M. Levitin, M. Miron, M. E. Snyder, T. Senda, J. Yuan, Y. L. Cheng, E. C. Bush, P. Dogra, P. Thapa, D. L. Farber, P. A. Sims, Single-cell transcriptomics of human T cells reveals tissue and activation signatures in health and disease. Nat. Commun. 10, 4706 (2019).

78. D. Morgan, V. Tergaonkar, Unraveling B cell trajectories at single cell resolution. Trends Immunol. 43, 210–229 (2022).

79. M. Hamada, Y. Tsunakawa, H. Jeon, M. K. Yadav, S. Takahashi, Role of MafB in macrophages. Exp. Anim. 69, 1–10 (2020).

80. L. M. Kelly, U. Englmeier, I. Lafon, M. H. Sieweke, T. Graf, MafB is an inducer of monocytic differentiation. EMBO J. 19, 1987–1997 (2000).

81. V. D. Cuevas, L. Anta, R. Samaniego, E. Orta-Zavalza, J. Vladimir de la Rosa, G. Baujat, Á. Domínguez-Soto, P. Sánchez-Mateos, M. M. Escribese, A. Castrillo, V. Cormier-Daire, M. A. Vega, Á. L. Corbí, MAFB Determines Human Macrophage Anti-Inflammatory Polarization: Relevance for the Pathogenic Mechanisms Operating in Multicentric Carpotarsal Osteolysis. J. Immunol. 198, 2070–2081 (2017).

82. K. Kang, S. H. Park, J. Chen, Y. Qiao, E. Giannopoulou, K. Berg, A. Hanidu, J. Li, G. Nabozny, K. Kang, K.-H. Park-Min, L. B. Ivashkiv, Interferon-γ Represses M2 Gene Expression in Human Macrophages by Disassembling Enhancers Bound by the Transcription Factor MAF. Immunity 47, 235–250.e4 (2017).

83. J. Fleischer, E. Soeth, N. Reiling, E. Grage-Griebenow, H. D. Flad, M. Ernst, Differential expression and function of CD80 (B7-1) and CD86 (B7-2) on human peripheral blood monocytes. Immunology 89, 592–598 (1996).

84. H. Fujita, Y. Hashimoto, S. Russell, B. Zieger, J. Ware, In Vivo Expression of Murine Platelet Glycoprotein Ibα. Blood 92, 488–495 (1998).

85. Q. Kimmerlin, C. Strassel, A. Eckly, F. Lanza, The tubulin code in platelet biogenesis. Semin. Cell Dev. Biol. 137, 63–73 (2023).

86. G. Varricchi, A. Pecoraro, S. Loffredo, R. Poto, F. Rivellese, A. Genovese, G. Marone, G. Spadaro, Heterogeneity of Human Mast Cells With Respect to MRGPRX2 Receptor Expression and Function. Front. Cell. Neurosci. 13, 299 (2019).

87. G. Cruse, D. Kaur, M. Leyland, P. Bradding, A novel FcεRIβ-chain truncation regulates human mast cell proliferation and survival. FASEB J. 24, 4047–4057 (2010).

88. K. R. Acharya, S. J. Ackerman, Eosinophil granule proteins: form and function. J. Biol. Chem. 289, 17406–17415 (2014).

89. J. Tavernier, R. Devos, S. Cornelis, T. Tuypens, J. Van der Heyden, W. Fiers, G. Plaetinck, A human high affinity interleukin-5 receptor (IL5R) is composed of an IL5-specific alpha chain and a beta chain shared with the receptor for GM-CSF. Cell 66, 1175–1184 (1991).

90. B. L. Cantarel, I. Korf, S. M. C. Robb, G. Parra, E. Ross, B. Moore, C. Holt, A. Sánchez Alvarado, M. Yandell, MAKER: an easy-to-use annotation pipeline designed for emerging model organism genomes. Genome Res. 18, 188–196 (2008).

91. M. Stanke, O. Keller, I. Gunduz, A. Hayes, S. Waack, B. Morgenstern, AUGUSTUS: ab initio prediction of alternative transcripts. Nucleic Acids Res. 34, W435–9 (2006).

92. I. Korf, Gene finding in novel genomes. BMC Bioinformatics 5, 59 (2004).

93. A. Shumate, S. L. Salzberg, Liftoff: accurate mapping of gene annotations. Bioinformatics 37, 1639–1643 (2021).

94. K. Katoh, K. Misawa, K.-I. Kuma, T. Miyata, MAFFT: a novel method for rapid multiple sequence alignment based on fast Fourier transform. Nucleic Acids Res. 30, 3059–3066 (2002).

95. A. Stamatakis, RAxML version 8: a tool for phylogenetic analysis and post-analysis of large phylogenies. Bioinformatics 30, 1312–1313 (2014).

96. S. Kumar, G. Stecher, M. Li, C. Knyaz, K. Tamura, MEGA X: Molecular Evolutionary Genetics Analysis across Computing Platforms. Mol. Biol. Evol. 35, 1547–1549 (2018).

97. Rambaut, A. Figtree v1.4.4. Institute of Evolutionary Biology, University of Edinburgh, Edinburgh. (2018).

98. S. R. Amend, K. C. Valkenburg, K. J. Pienta, Murine Hind Limb Long Bone Dissection and Bone Marrow Isolation. J. Vis. Exp., doi: 10.3791/53936 (2016).

99. A. M. Bolger, M. Lohse, B. Usadel, Trimmomatic: a flexible trimmer for Illumina sequence data. Bioinformatics 30, 2114–2120 (2014).

100. A. Dobin, C. A. Davis, F. Schlesinger, J. Drenkow, C. Zaleski, S. Jha, P. Batut, M. Chaisson, T. R. Gingeras, STAR: ultrafast universal RNA-seq aligner. Bioinformatics 29, 15–21 (2013).

101. Y. Liao, G. K. Smyth, W. Shi, featureCounts: an efficient general purpose program for assigning sequence reads to genomic features. Bioinformatics 30, 923–930 (2014).

102. B. Langmead, S. L. Salzberg, Fast gapped-read alignment with Bowtie 2. Nat. Methods 9, 357–359 (2012).

103. Y. Zhang, T. Liu, C. A. Meyer, J. Eeckhoute, D. S. Johnson, B. E. Bernstein, C. Nusbaum, R. M. Myers, M. Brown, W. Li, X. S. Liu, Model-based analysis of ChIP-Seq (MACS). Genome Biol. 9, R137 (2008).

104. Q. Li, J. B. Brown, H. Huang, P. J. Bickel, Measuring reproducibility of high-throughput experiments. aoas 5, 1752–1779 (2011).

105. J. T. Robinson, H. Thorvaldsdóttir, W. Winckler, M. Guttman, E. S. Lander, G. Getz, J. P. Mesirov, Integrative genomics viewer. Nat. Biotechnol. 29, 24–26 (2011).

106. E. Slobodyanyuk, C. Cattoglio, T.-H. S. Hsieh, Mapping Mammalian 3D Genomes by Micro-C. Methods Mol. Biol. 2532, 51–71 (2022).

107. T.-H. S. Hsieh, C. Cattoglio, E. Slobodyanyuk, A. S. Hansen, O. J. Rando, R. Tjian, X. Darzacq, Resolving the 3D Landscape of Transcription-Linked Mammalian Chromatin Folding. Mol. Cell 78, 539–553.e8 (2020).

108. P. J. Batut, X. Y. Bing, Z. Sisco, J. Raimundo, M. Levo, M. S. Levine, Genome organization controls transcriptional dynamics during development. Science 375, 566–570 (2022).

109. X. Bing, W. Ke, M. Fujioka, A. Kurbidaeva, S. Levitt, M. Levine, P. Schedl, J. B. Jaynes, Chromosome Structure I: Loop extrusion or boundary:boundary pairing?, bioRxiv (2024)p. 2023.11.17.567501.

110. W. Ke, M. Fujioka, P. Schedl, J. B. Jaynes, Chromosome Structure II: Stem-loops and Circle-loops, bioRxiv (2024)p. 2024.01.02.573928.

111. H. Li, R. Durbin, Fast and accurate short read alignment with Burrows-Wheeler transform. Bioinformatics 25, 1754–1760 (2009).

112. N. Abdennur, L. A. Mirny, Cooler: scalable storage for Hi-C data and other genomically labeled arrays. Bioinformatics 36, 311–316 (2020).

113. P. Kerpedjiev, N. Abdennur, F. Lekschas, C. McCallum, K. Dinkla, H. Strobelt, J. M. Luber, S. B. Ouellette, A. Azhir, N. Kumar, J. Hwang, S. Lee, B. H. Alver, H. Pfister, L. A. Mirny, P. J. Park, N. Gehlenborg, HiGlass: web-based visual exploration and analysis of genome interaction maps. Genome Biol. 19, 125 (2018).

114. C. E. Grant, T. L. Bailey, W. S. Noble, FIMO: scanning for occurrences of a given motif. Bioinformatics 27, 1017–1018 (2011).

115. Z. Zou, T. Ohta, F. Miura, S. Oki, ChIP-Atlas 2021 update: a data-mining suite for exploring epigenomic landscapes by fully integrating ChIP-seq, ATAC-seq and Bisulfite-seq data. Nucleic Acids Res. 50, W175–W182 (2022).

116. Z. Zou, T. Ohta, S. Oki, ChIP-Atlas 3.0: a data-mining suite to explore chromosome architecture together with large-scale regulome data. Nucleic Acids Res., doi: 10.1093/nar/gkae358 (2024).

117. S. Oki, T. Ohta, G. Shioi, H. Hatanaka, O. Ogasawara, Y. Okuda, H. Kawaji, R. Nakaki, J. Sese, C. Meno, ChIP-Atlas: a data-mining suite powered by full integration of public ChIP-seq data. EMBO Rep. 19, e46255 (2018).

118. A. Lamiable, P. Thévenet, J. Rey, M. Vavrusa, P. Derreumaux, P. Tufféry, PEP-FOLD3: faster de novo structure prediction for linear peptides in solution and in complex. Nucleic Acids Res. 44, W449–54 (2016).

119. I. Wiegand, K. Hilpert, R. E. W. Hancock, Agar and broth dilution methods to determine the minimal inhibitory concentration (MIC) of antimicrobial substances. Nat. Protoc. 3, 163–175 (2008).

120. P. Stothard, The sequence manipulation suite: JavaScript programs for analyzing and formatting protein and DNA sequences. Biotechniques 28, 1102, 1104 (2000).

121. B. D. Rosen, D. M. Bickhart, R. D. Schnabel, S. Koren, C. G. Elsik, E. Tseng, T. N. Rowan, W. Y. Low, A. Zimin, C. Couldrey, R. Hall, W. Li, A. Rhie, J. Ghurye, S. D. McKay, F. Thibaud-Nissen, J. Hoffman, B. M. Murdoch, W. M. Snelling, T. G. McDaneld, J. A. Hammond, J. C. Schwartz, W. Nandolo, D. E. Hagen, C. Dreischer, S. J. Schultheiss, S. G. Schroeder, A. M. Phillippy, J. B. Cole, C. P. Van Tassell, G. Liu, T. P. L. Smith, J. F. Medrano, De novo assembly of the cattle reference genome with single-molecule sequencing. Gigascience 9 (2020).

122. B. P. Bentley, T. Carrasco-Valenzuela, E. K. S. Ramos, H. Pawar, L. Souza Arantes, A. Alexander, S. M. Banerjee, P. Masterson, M. Kuhlwilm, M. Pippel, J. Mountcastle, B. Haase, M. Uliano-Silva, G. Formenti, K. Howe, W. Chow, A. Tracey, Y. Sims, S. Pelan, J. Wood, K. Yetsko, J. R. Perrault, K. Stewart, S. R. Benson, Y. Levy, E. V. Todd, H. B. Shaffer, P. Scott, B. T. Henen, R. W. Murphy, D. W. Mohr, A. F. Scott, D. J. Duffy, N. J. Gemmell, A. Suh, S. Winkler, F. Thibaud-Nissen, M. F. Nery, T. Marques-Bonet, A. Antunes, Y. Tikochinski, P. H. Dutton, O. Fedrigo, E. W. Myers, E. D. Jarvis, C. J. Mazzoni, L. M. Komoroske, Divergent sensory and immune gene evolution in sea turtles with contrasting demographic and life histories. Proc. Natl. Acad. Sci. U. S. A. 120, e2201076120 (2023).

123. T. S. Kalbfleisch, E. S. Rice, M. S. DePriest Jr, B. P. Walenz, M. S. Hestand, J. R. Vermeesch, B. L. O Connell, I. T. Fiddes, A. O. Vershinina, N. F. Saremi, J. L. Petersen, C. J. Finno, R. R. Bellone, M. E. McCue, S. A. Brooks, E. Bailey, L. Orlando, R. E. Green, D. C. Miller, D. F. Antczak, J. N. MacLeod, Improved reference genome for the domestic horse increases assembly contiguity and composition. Commun Biol 1, 197 (2018).

124. V. C. Mason, G. Li, P. Minx, J. Schmitz, G. Churakov, L. Doronina, A. D. Melin, N. J. Dominy, N. T.-L. Lim, M. S. Springer, R. K. Wilson, W. C. Warren, K. M. Helgen, W. J. Murphy, Genomic analysis reveals hidden biodiversity within colugos, the sister group to primates. Sci Adv 2, e1600633 (2016).

125. V. A. Schneider, T. Graves-Lindsay, K. Howe, N. Bouk, H.-C. Chen, P. A. Kitts, T. D. Murphy, K. D. Pruitt, F. Thibaud-Nissen, D. Albracht, R. S. Fulton, M. Kremitzki, V. Magrini, C. Markovic, S. McGrath, K. M. Steinberg, K. Auger, W. Chow, J. Collins, G. Harden, T. Hubbard, S. Pelan, J. T. Simpson, G. Threadgold, J. Torrance, J. M. Wood, L. Clarke, S. Koren, M. Boitano, P. Peluso, H. Li, C.-S. Chin, A. M. Phillippy, R. Durbin, R. K. Wilson, P. Flicek, E. E. Eichler, D. M. Church, Evaluation of GRCh38 and de novo haploid genome assemblies demonstrates the enduring quality of the reference assembly. Genome Res. 27, 849–864 (2017).

126. J.-Y. Hu, Z.-Q. Hao, L. Frantz, S.-F. Wu, W. Chen, Y.-F. Jiang, H. Wu, W.-M. Kuang, H. Li, Y.-P. Zhang, L. Yu, Genomic consequences of population decline in critically endangered pangolins and their demographic histories. Natl Sci Rev 7, 798–814 (2020).

127. T. S. Mikkelsen, M. J. Wakefield, B. Aken, C. T. Amemiya, J. L. Chang, S. Duke, M. Garber, A. J. Gentles, L. Goodstadt, A. Heger, J. Jurka, M. Kamal, E. Mauceli, S. M. J. Searle, T. Sharpe, M. L. Baker, M. A. Batzer, P. V. Benos, K. Belov, M. Clamp, A. Cook, J. Cuff, R. Das, L. Davidow, J. E. Deakin, M. J. Fazzari, J. L. Glass, M. Grabherr, J. M. Greally, W. Gu, T. A. Hore, G. A. Huttley, M. Kleber, R. L. Jirtle, E. Koina, J. T. Lee, S. Mahony, M. A. Marra, R. D. Miller, R. D. Nicholls, M. Oda, A. T. Papenfuss, Z. E. Parra, D. D. Pollock, D. A. Ray, J. E. Schein, T. P. Speed, K. Thompson, J. L. VandeBerg, C. M. Wade, J. A. Walker, P. D. Waters, C. Webber, J. R. Weidman, X. Xie, M. C. Zody, Broad Institute Genome Sequencing Platform, Broad Institute Whole Genome Assembly Team, J. A. M. Graves, C. P. Ponting, M. Breen, P. B. Samollow, E. S. Lander, K. Lindblad-Toh, Genome of the marsupial Monodelphis domestica reveals innovation in non-coding sequences. Nature 447, 167–177 (2007).

128. Y. Zhou, L. Shearwin-Whyatt, J. Li, Z. Song, T. Hayakawa, D. Stevens, J. C. Fenelon, E. Peel, Y. Cheng, F. Pajpach, N. Bradley, H. Suzuki, M. Nikaido, J. Damas, T. Daish, T. Perry, Z. Zhu, Y. Geng, A. Rhie, Y. Sims, J. Wood, B. Haase, J. Mountcastle, O. Fedrigo, Q. Li, H. Yang, J. Wang, S. D. Johnston, A. M. Phillippy, K. Howe, E. D. Jarvis, O. A. Ryder, H. Kaessmann, P. Donnelly, J. Korlach, H. A. Lewin, J. Graves, K. Belov, M. B. Renfree, F. Grutzner, Q. Zhou, G. Zhang, Platypus and echidna genomes reveal mammalian biology and evolution. Nature 592, 756–762 (2021).

129. R. N. Johnson, D. O’Meally, Z. Chen, G. J. Etherington, S. Y. W. Ho, W. J. Nash, C. E. Grueber, Y. Cheng, C. M. Whittington, S. Dennison, E. Peel, W. Haerty, R. J. O’Neill, D. Colgan, T. L. Russell, D. E. Alquezar-Planas, V. Attenbrow, J. G. Bragg, P. A. Brandies, A. Y.-Y. Chong, J. E. Deakin, F. Di Palma, Z. Duda, M. D. B. Eldridge, K. M. Ewart, C. J. Hogg, G. J. Frankham, A. Georges, A. K. Gillett, M. Govendir, A. D. Greenwood, T. Hayakawa, K. M. Helgen, M. Hobbs, C. E. Holleley, T. N. Heider, E. A. Jones, A. King, D. Madden, J. A. M. Graves, K. M. Morris, L. E. Neaves, H. R. Patel, A. Polkinghorne, M. B. Renfree, C. Robin, R. Salinas, K. Tsangaras, P. D. Waters, S. A. Waters, B. Wright, M. R. Wilkins, P. Timms, K. Belov, Adaptation and conservation insights from the koala genome. Nat. Genet. 50, 1102–1111 (2018).

130. D. Jebb, Z. Huang, M. Pippel, G. M. Hughes, K. Lavrichenko, P. Devanna, S. Winkler, L. S. Jermiin, E. C. Skirmuntt, A. Katzourakis, L. Burkitt-Gray, D. A. Ray, K. A. M. Sullivan, J. G. Roscito, B. M. Kirilenko, L. M. Dávalos, A. P. Corthals, M. L. Power, G. Jones, R. D. Ransome, D. K. N. Dechmann, A. G. Locatelli, S. J. Puechmaille, O. Fedrigo, E. D. Jarvis, M. Hiller, S. C. Vernes, E. W. Myers, E. C. Teeling, Six reference-quality genomes reveal evolution of bat adaptations. Nature 583, 578–584 (2020).

131. P. Andrade, C. Pinho, G. Pérez I de Lanuza, S. Afonso, J. Brejcha, C.-J. Rubin, O. Wallerman, P. Pereira, S. J. Sabatino, A. Bellati, D. Pellitteri-Rosa, Z. Bosakova, I. Bunikis, M. A. Carretero, N. Feiner, P. Marsik, F. Paupério, D. Salvi, L. Soler, G. M. While, T. Uller, E. Font, L. Andersson, M. Carneiro, Regulatory changes in pterin and carotenoid genes underlie balanced color polymorphisms in the wall lizard. Proc. Natl. Acad. Sci. U. S. A. 116, 5633–5642 (2019).

132. M. R. Stammnitz, K. Gori, Y. M. Kwon, E. Harry, F. J. Martin, K. Billis, Y. Cheng, A. Baez-Ortega, W. Chow, S. Comte, H. Eggertsson, S. Fox, R. Hamede, M. Jones, B. Lazenby, S. Peck, R. Pye, M. A. Quail, K. Swift, J. Wang, J. Wood, K. Howe, M. R. Stratton, Z. Ning, E. P. Murchison, The evolution of two transmissible cancers in Tasmanian devils. Science 380, 283–293 (2023).

133. A. D. Foote, Y. Liu, G. W. C. Thomas, T. Vinař, J. Alföldi, J. Deng, S. Dugan, C. E. van Elk, M. E. Hunter, V. Joshi, Z. Khan, C. Kovar, S. L. Lee, K. Lindblad-Toh, A. Mancia, R. Nielsen, X. Qin, J. Qu, B. J. Raney, N. Vijay, J. B. W. Wolf, M. W. Hahn, D. M. Muzny, K. C. Worley, M. T. P. Gilbert, R. A. Gibbs, Convergent evolution of the genomes of marine mammals. Nat. Genet. 47, 272–275 (2015).

134. Y. Fan, Z.-Y. Huang, C.-C. Cao, C.-S. Chen, Y.-X. Chen, D.-D. Fan, J. He, H.-L. Hou, L. Hu, X.-T. Hu, X.-T. Jiang, R. Lai, Y.-S. Lang, B. Liang, S.-G. Liao, D. Mu, Y.-Y. Ma, Y.-Y. Niu, X.-Q. Sun, J.-Q. Xia, J. Xiao, Z.-Q. Xiong, L. Xu, L. Yang, Y. Zhang, W. Zhao, X.-D. Zhao, Y.-T. Zheng, J.-M. Zhou, Y.-B. Zhu, G.-J. Zhang, J. Wang, Y.-G. Yao, Genome of the Chinese tree shrew. Nat. Commun. 4, 1426 (2013).

135. T. Mitros, J. B. Lyons, A. M. Session, J. Jenkins, S. Shu, T. Kwon, M. Lane, C. Ng, T. C. Grammer, M. K. Khokha, J. Grimwood, J. Schmutz, R. M. Harland, D. S. Rokhsar, A chromosome-scale genome assembly and dense genetic map for Xenopus tropicalis. Dev. Biol. 452, 8–20 (2019).

